# Cardiac glycosides restore autophagy flux in an iPSC-derived neuronal model of WDR45 deficiency

**DOI:** 10.1101/2023.09.13.556416

**Authors:** Apostolos Papandreou, Nivedita Singh, Lorita Gianfrancesco, Dimitri Budinger, Katy Barwick, Alexander Agrotis, Christin Luft, Ying Shao, An-Sofie Lenaerts, Allison Gregory, Suh Young Jeong, Penelope Hogarth, Susan Hayflick, Serena Barral, Janos Kriston-Vizi, Paul Gissen, Manju A Kurian, Robin Ketteler

## Abstract

Beta-Propeller Protein-Associated Neurodegeneration (BPAN) is one of the commonest forms of Neurodegeneration with Brain Iron Accumulation, caused by mutations in the gene encoding the autophagy-related protein, WDR45. The mechanisms linking autophagy, iron overload and neurodegeneration in BPAN are poorly understood and, as a result, there are currently no disease-modifying treatments for this progressive disorder. We have developed a patient-derived, induced pluripotent stem cell (iPSC)-based midbrain dopaminergic neuronal cell model of BPAN (3 patient, 2 age-matched controls and 2 isogenic control lines) which shows defective autophagy and aberrant gene expression in key neurodegenerative, neurodevelopmental and collagen pathways. A high content imaging-based medium-throughput blinded drug screen using the FDA-approved Prestwick library identified 5 cardiac glycosides that both corrected disease-related defective autophagosome formation and restored BPAN-specific gene expression profiles. Our findings have clear translational potential and emphasise the utility of iPSC-based modelling in elucidating disease pathophysiology and identifying targeted therapeutics for early-onset monogenic disorders.

## Introduction

Neurodegeneration with Brain Iron Accumulation (NBIA) encompasses a heterogeneous group of neurodegenerative conditions associated with excess iron accumulating in the brain.^1, 2^ Beta-Propeller Protein-Associated Neurodegeneration (BPAN) is one of the more recently described NBIA disorders. BPAN is caused by mutations in *WDR45* (WD repeat domain 45),^3, 4, 5^ located on Xp11.23 and also known as *WIPI4* (WD repeat domain phosphoinositide-interacting protein 4).^5, 6^ The encoded protein has a role in the early stages of autophagy, a cellular innate homeostatic process whereby damaged or redundant cargo (*e.g.,* protein aggregates, damaged organelles) is engulfed in a double lipid bilayer membrane structure (the autophagosome) in response to various stresses and/or stimuli and delivered to the lysosome for degradation and recycling.^5, 7, 8^ Clinically, BPAN is typically characterised by two distinct phases: a complex but relatively static neurodevelopmental phenotype in infancy and childhood with global developmental delay, seizures, disrupted sleep pattern and behavioural features (including stereotypies and hyperphagia), followed by neurological regression with dementia and prominent parkinsonism in adolescence or early adulthood. Post-mortem analysis confirms neuropathological changes in the entire brain, especially within the substantia nigra and globus pallidus, with iron accumulation, neuronal loss, and reactive gliosis, as well as excess hyperphosphorylated tau deposition.^9^ BPAN is a devastating early-onset neurodegenerative condition associated with severe morbidity and high risk of premature mortality, for which there are currently no disease-modifying treatments. For BPAN, the links between autophagy, iron metabolism and neurodegeneration are poorly understood. Autophagy is a highly conserved intracellular trafficking pathway that is important for cellular homeostasis, particularly for post-mitotic and metabolically active tissues, such as neurons^10^. Dysregulation of autophagy is associated with numerous diseases including ageing, cancer and neurodegeneration.^11, 12, 13^ Autophagosome formation is a multi-step process that is initiated by the serine/threonine kinase Unc-51-like kinase 1/2 (ULK1/2) and the phosphoinositide kinase vps34/PI3K class III. These kinases then lead to the formation of the autophagosomal membrane, through production of phosphatidylinositol 3-phosphate (PI3P), a series of ubiquitin-like conjugation reactions and the contribution of PI3P-effector proteins such as the WD-repeat proteins interacting with phosphoinositides (WIPI).^8, 14^ WIPI subunits in mammals associate with autophagosomal membranes through a phosphoinositide-binding Phe-Arg-Arg-Gly motif (FRRG) within a seven–β propeller structure.^15, 16, 17^ WDR45 (WIPI4) is essential for the early stages of autophagosome formation.^18, 19^ In *C. elegans* studies,^18^ the WIPI4 homologue EPG6 forms a complex with Autophagy-Related Protein 2 (ATG2), another autophagy-related protein, and regulates the generation of autophagosomes from omegasomes. Accumulation of (abnormally enlarged) Double FYVE-containing protein 1 (DFCP1)-labelled omegasomes was seen in *WDR45*-Knockdown Normal Rat Kidney cells^18^ whereas, upon starvation (a known autophagy flux activator), accumulation of Microtubule-associated protein 1A/1B-light chain 3 (LC3)-labelled autophagosomes and Autophagy-Related Protein 5 (ATG5) puncta was also seen in *C. Elegans*-based studies. ATG5 dissociates from the autophagosomal membrane upon the autophagosome’s completion in *S. cerevisiae* models.^20^ The evidence thus suggests that *WDR45* knockdown leads to defective autophagosome formation.^18^ Similarly, studies in human lymphoblastoid cell lines derived from patients with *WDR45* mutations showed accumulation of LC3-marked autophagosomes compared to controls, both in basal conditions and after autophagy flux induction^5^. These LC3-positive structures co-localised with ATG9A, which is absent from completed autophagosomes, again suggesting that the autophagosome formation in WDR45 deficiency is abnormal or impaired.^5^ Human Bone Osteosarcoma Epithelial (U2OS) cell studies also confirmed that WDR45 forms a complex with ATG2A through two critical residues, N15 and D17, which are conserved in human WIPI proteins. This complex also contains AMP-activated protein kinase (AMPK) and ULK1 in fed conditions; however, upon starvation, and possibly due to AMPK activation, the WDR45/ATG2A complex is proposed to dissociate from AMPK and ULK1 and localise to the autophagosome, possibly supporting autophagosomal maturation.^7^

To date, BPAN disease modelling has provided some useful insights into disease pathophysiology. Central nervous system (CNS)-specific *WDR45* knockout mice displayed defects in autophagy-mediated clearance of protein aggregates and recapitulated several hallmarks of BPAN, such as cognitive impairment and defective axonal homeostasis^6^. Another study using constitutive *WDR45* knockout mice showed accumulation of endoplasmic reticulum (ER) stress in neurons over time, leading to neuronal apoptosis.^21^ Studies based on cell models ^22^ have shown reduced LC3 in BPAN fibroblasts and neurons compared to controls, as well as iron accumulation and possible mitochondrial and lysosomal dysfunction. Additionally, there is some indication that induction of autophagy in BPAN neurons might lead to reversal of iron accumulation or ER stress.^21, 22^ Overall, autophagy is likely to be a key dysfunctional pathway in BPAN; as such, overcoming the underlying autophagy defect may provide a translational path for future therapeutic strategies.

For BPAN, better understanding of the disease mechanisms and development of novel precision therapies constitutes an urgent and unmet research priority. In order to study BPAN in a physiologically relevant context, we have established a patient-derived induced pluripotent stem cell (iPSC)-based midbrain dopaminergic (mDA) neuronal model, which has revealed key disease-specific phenotypes. Moreover, a medium-throughput drug screen using high content imaging-based immunofluorescence assays has identified several compounds that ameliorate disease-specific phenotypes *in vitro,* with clear potential for clinical translation.

## Results

### Patient and control iPSCs efficiently differentiate into mature mDA neurons

IPSC lines were generated^23^ from dermal fibroblasts of BPAN patients with loss-of-function mutations in *WDR45* [Patient 01 (female): c.344+2T>A, p.Ile116Argfs*3; Patient 02 (male): c.19C>T, p.Arg7*; Patient 03 (female): c.700C>T, p.Arg234*]^24^. Control iPSCs were similarly generated from two age-matched healthy individuals. Two isogenic control lines were created by CRISPR-Cas9 correction of the c.19C>T and c.700C>T variants in patients 2 and 3, respectively (Supplementary Table 1).

All iPSC lines exhibited pluripotency (Supplementary Fig.1A-E).^25^ Patient-derived iPSC lines maintained their *WDR45* mutations (Supplementary Fig. 1F) and genomic integrity following reprogramming (Supplementary Fig. 1G), with successful correction of the mutation in the isogenic controls (Supplementary Fig. 1H).

iPSC lines were differentiated into mDA neurons using a modified dual SMAD inhibition protocol (Supplementary Fig. 2A).^26, 27^ After 11 days of differentiation, co-expression of LIM homeobox transcription factor 1 alpha (LMX1A) and forkhead box protein A2 (FOXA2) in a large percentage of cells in all lines signified effective initiation of differentiation into mDA neurons (Supplementary Fig. 2B-C). Comparable mDA progenitors were present in both patient and control lines, with typical midbrain precursor gene expression profiles (Supplementary Fig. 2D). After 65 days of differentiation, control and patient mDA neurons showed typical mature-derived morphology with comparable protein (Fig. 1A, 1B) and gene expression profiles for derived-maturity markers (Supplementary Fig. 2E), in line with published studies.^26, 27^

**Figure 1.**
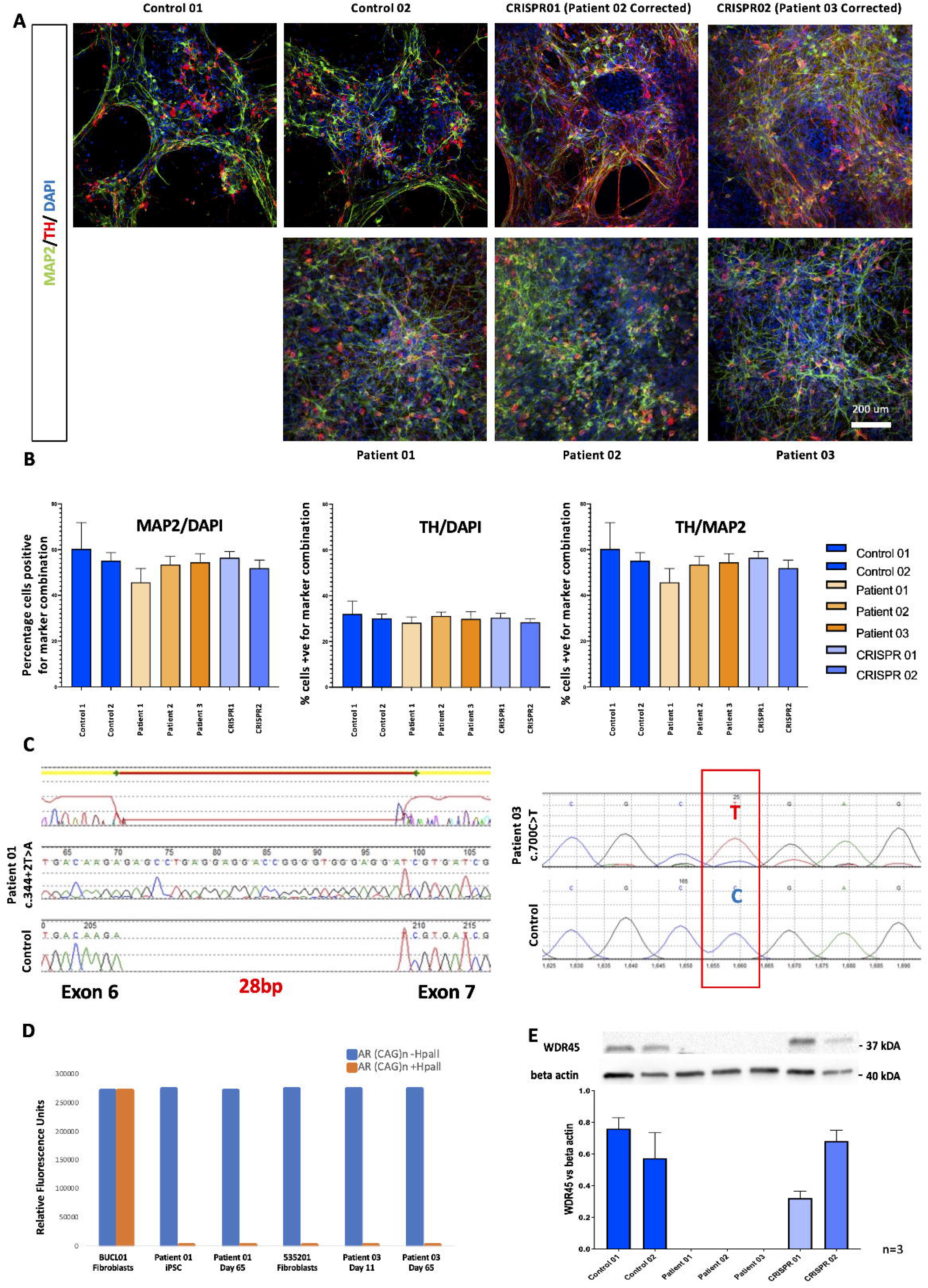
BPAN and control iPSC line differentiation into mDA neurons. **(A)** Immunofluorescence analysis of MAP2/TH in patient and control lines at day 65 of differentiation. Mature mDA neurons exhibit typical morphology and express pan-neuronal (MAP2) and mDA-specific (TH) markers. Nuclei were counterstained with DAPI. Scale bar, 200 μm. n=3 biological replicates per line. **(B)** Quantification of MAP2/TH abundance in control and patient-derived neurons (n=3 biological replicates per line, 3 individual images analysed per replicate). Percentages were calculated after manual counting of cells on ImageJ/Fiji (approximately 500 nuclei counted per image, followed by counting of cells also staining positive for TH and/or MAP2). **(C)** Chromatograms from WDR45 cDNA sequencing in female BPAN iPSC lines. In Patient 01, the cDNA has 28 additional bp, while for Patient 03, the c.700C>T leading to an early stop codon is retained. **(D)** Relative Fluorescence Units of Androgen Receptor CAG repeat PCR products from female BPAN lines, in the presence or absence of methylation-sensitive restriction enzyme Hpall. Patient 01 fibroblasts (BUCL01) exhibit a random XCI pattern, with two PCR bands detectable in the presence and absence of Hpall. For patient 03 (535-201), there is practically only one detectable band in fibroblasts (535-201), signifying a skewing of XCI towards the expression of only one allele. For both patients, iPSC lines and derived neurons have very skewed patterns of XCI, with PCR bands practically undetectable in the presence of Hpall. n=1 biological replicate per line. **(E)** Cropped immunoblot of total WDR45 and beta actin protein expression at Day 65, and quantification of WDR45 relative to the loading control. n=3 biological replicates for each line. Error bars represent the Standard Error of Mean. Statistics were calculated using ANOVA.

### Patient-derived mDA neurons express mutant *WDR45* allele with absent WDR45 protein

Since X Chromosome Inactivation (XCI) can influence the modelling of X-linked disorders, *WDR45* allele expression was examined for both female patient lines (Patients 01 and 03) in fibroblasts, iPSCs and mDA neurons. Although XCI is likely maintained throughout reprogramming and differentiation,^28, 29, 30^ XCI erosion leading to expression of the wild-type allele from the previously inactive X can occur.^31^ Differences in methylation at the X-linked Androgen Receptor locus (examined via fluorescent PCR in the presence and absence of the methylation-sensitive restriction enzyme HpaII)^32^ confirmed that only one *WDR45* allele was expressed in iPSCs and neurons derived from female Patient 01 and Patient 03. cDNA sequencing confirmed expression of the mutant *WDR45* allele at iPSC stage (Fig. 1C and 1D). Moreover, WDR45 protein was almost undetectable in all 3 patient-derived mDA progenitor cultures at Day 11 (Supplementary Fig. 2F), and also at day 65, of differentiation (Fig. 1E). These results indicate that XCI patterns in both female lines remained skewed towards the expression of the mutated WDR45 allele during neuronal differentiation.

### Mature-derived mDA cultures show disease-specific dysregulated gene expression profiles in key neurodevelopmental and neurodegenerative pathways

Bulk RNA sequencing (RNAseq) at Day 65 mDA cultures^33^ identified 59 differentially expressed genes (28 overexpressed, 31 underexpressed) in BPAN lines, when compared to controls. Subsequent Gene Ontology (GO) term annotation^34, 35, 36^ and Kyoto Encyclopedia of Genes and Genomes (KEGG) pathway^37, 38, 39^ enrichment analysis^40^ revealed disease-specific defects in several intracellular pathways, including those involved in CNS development (KEGG:hsa04722, neurotrophin signalling pathway,^41^ p=0.02; *ARHGDIG, MAPK7*), ferroptosis (KEGG:hsa04216, p=0.02; *ACSL6, SLC40A1*),^42^ and collagen metabolism (e.g. GO:0030574, p=4.1×10^−11^; *COL1A1, COL18A1, COL3A1, COL1A2, COL5A1, COL4A1, MMP2, COL5A2*) (Fig. 2A-C; Supplementary Fig. 3A-B), affecting several collagen pathways. Comparison of gene expression profiles in Patients 02 and 03 versus the corresponding isogenic controls (CRISPR 01 and 02, respectively) showed similarly affected pathways in patient-derived neurons (Supplementary Fig. 3C-D).

**Figure 2.**
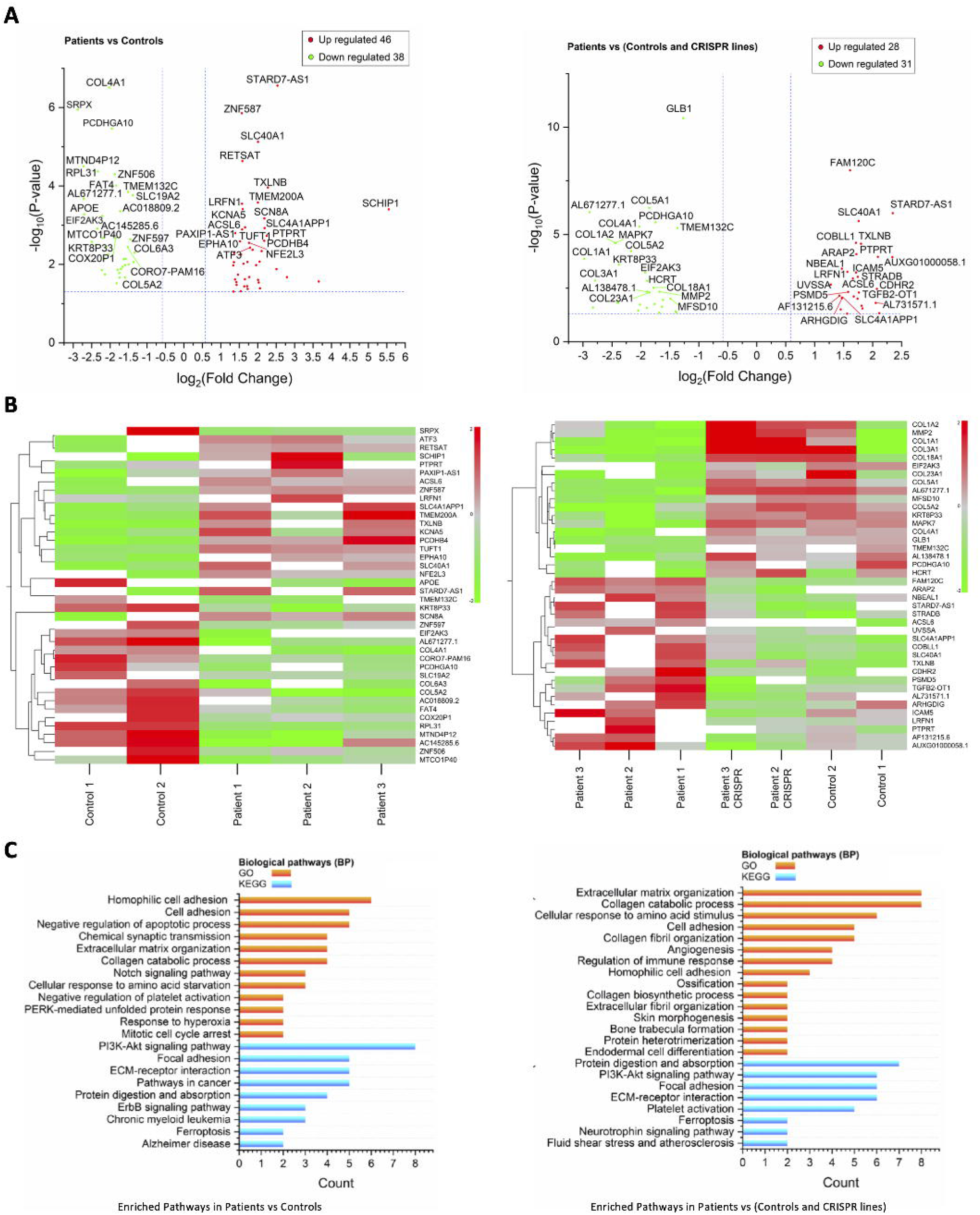
RNASeq at Day 65 of differentiation. **(A)** Volcano plots comparing gene expression profiles patient lines versus controls. Overexpressed genes are shown on the right of the X axis while underexpressed genes of the left. **(B)** Heat maps depicting gene underexpression and overexpression in gradients of green and red, respectively. **(C)** GO Term and KEGG pathway enrichment analysis depicting intracellular pathways affected in BPAN. Pathways with the most significant Fold Enrichment (Y axis) are shown; differentially expressed gene count is depicted on the X axis. n=3 biological replicates for all lines, analysis based on median TPM values. P-values of <0.05 and fold change of <0.5 or >2 [-1<log2(FC)>1] (Student’s t-test) were set as statistically significant cut-offs. The top 40 genes (as per lowest p-values) are labelled in volcano plots and heat maps. ATG= autophagy-related gene, TPM= transcript per million.

In the larger analysis including all lines, we also observed several differentially expressed genes involved in neuronal development or homeostasis maintenance. Firstly, *PCDHGA1*0, from the family of protocadherins with a role in neurogenesis and neural network development,^43^ and linked to intellectual disability.^44^ *MAPK7*, involved in neural development,^45, 46^ was under-expressed in patient mDA neurons. Under-expression of *EIF2AK3*, with a key role in the ER stress-related cellular response and apoptosis,^47^ was also observed. Another notably under-expressed gene was *GLB1* (related to GM1 gangliosidosis,^48^ a childhood-onset neurodegenerative condition that can clinically and radiologically mimic NBIA).^49, 50^ Finally, *HCRT* was also under-expressed, encoding a precursor for orexins, that in turn regulate sleep, arousal and feeding behaviour.^51, 52^ Conversely, overexpressed genes in BPAN neurons included *SLC40A1* (involved in ferroptosis).^42^ Overexpression of *FAM120C*, which has been linked to autism spectrum disorder,^44^ was also seen. *PTPRT* and *LRFN1*, two genes linked to neuronal development and synaptic formation,^53, 54^ were also overexpressed in BPAN lines. Additionally, overexpression of *ICAM5,* implicated in the pathogenesis of Fragile X syndrome^55^ was seen in patient mDA neurons. Finally, *ALCS6*, which exerts neuroprotection in the CNS via enrichment of Docosahexanoic acid,^56^ was overexpressed in BPAN compared to control and CRISPR-corrected lines.

### BPAN lines exhibit abnormal autophagy flux

We also established high content imaging-based immunofluorescence assays to examine autophagy flux in BPAN lines. We designed an assay to examine LC3 puncta formation, representative of autophagosomes. Autophagy is a very dynamic process; therefore, to minimise fluctuations in autophagy flux due to unaccounted factors, all patient and control lines were cultured on the same 96-well plates, at the same seeding density, and treated concurrently for 3 hours with autophagy flux inducers and/ or inhibitors. Treatments consisted of DMSO (negative control), bafilomycin A1 (autophagy flux blocker), and Torin 1 (autophagy inducer).^57, 58, 59^ Afterwards, cells were fixed, stained, and imaged.

We initially observed that BPAN fibroblasts exhibited defective autophagy flux, with fewer LC3 puncta forming in all conditions when compared to the age-ma*tched control fibroblasts (Supplementary Fig. 4A, 4B). Significantly fewer LC3 puncta were also observed in Day 11 patient-derived mDA progenitors in basal (DMSO-treated) conditions when compared to age-matched and CRISPR-corrected controls (Fig. 3A, Supplementary Figure 4C). Comparison between Control 02 and Patient 03 did not reach statistical significance using the two-tailed t-test (p=0.051); however, the corresponding isogenic control line CRISPR02 produced significantly higher LC3 puncta numbers than Patient 02. Additionally, the LC3 puncta production defect was reversible in all patient lines with autophagy flux induction by Torin 1 administration (Fig. 3B). Finally, BPAN Day 11 mDA progenitors exhibited similar defects in LC3 puncta formation after autophagy flux induction or inhibition (Supplementary Fig. 4D, 4E).

**Figure 3.**
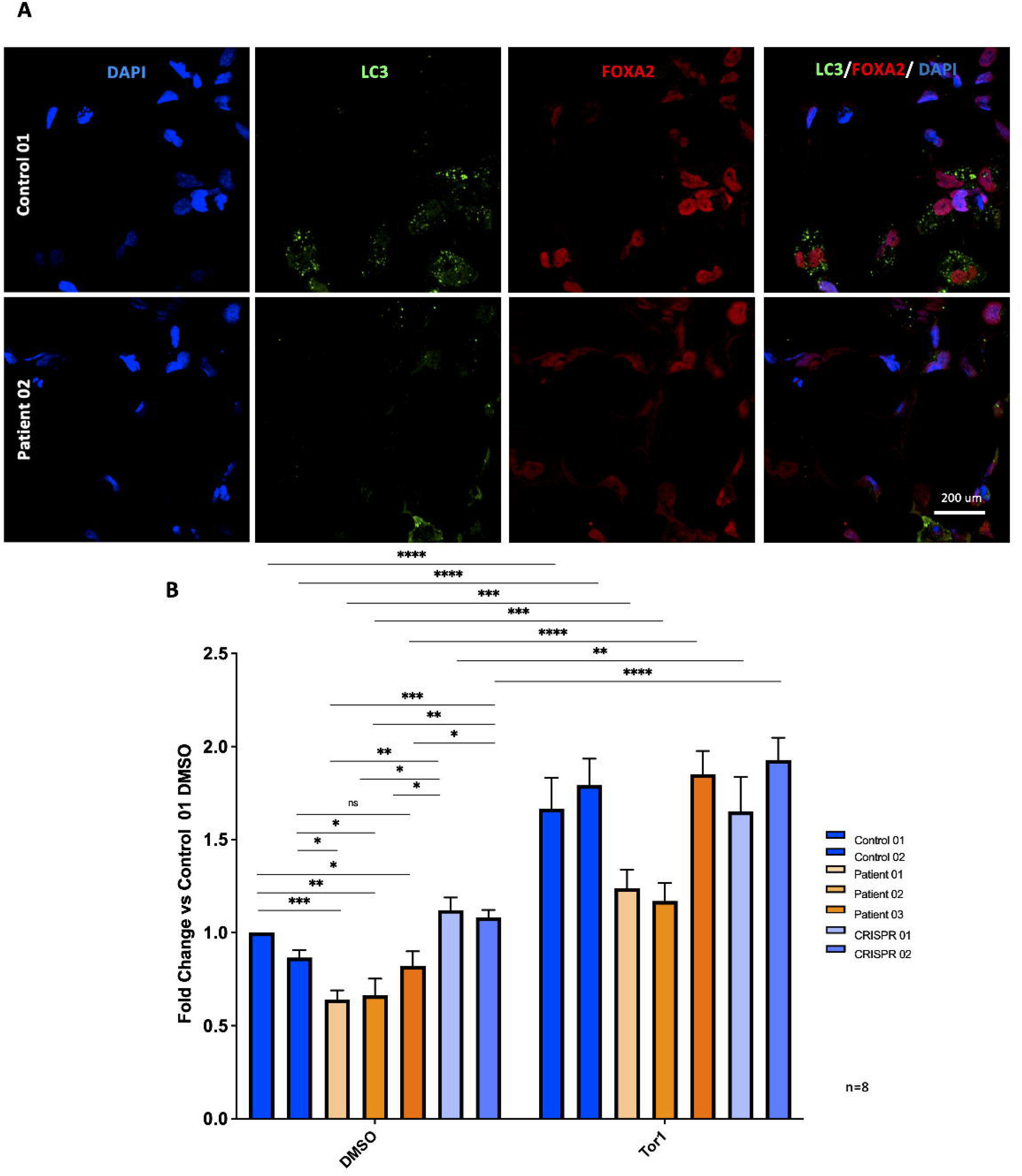
High content imaging LC3 assay for drug screening. **(A)** High Content Imaging immunofluorescence analysis of LC3/ FOXA2 in patient and control ventral midbrain progenitors at day 11 of differentiation. Representative images. Cell density at 15,000 cells/well. n=8 independent differentiations/ biological replicates for each line. **(B)** Quantification of LC3 puncta/ nuclei in control and patient-derived neurons DMSO and Torin 1 treatments were of 3-hour durations. n=8 independent differentiations/ biological replicates for each line, each condition per line tested in technical duplicates, 20 fields imaged per well. For statistical analysis, the Student’s unpaired two tailed t-test was used. Error bars represent the Standard Error of Mean.

### Medium throughput screening identifies therapeutic candidates that induce autophagy flux and reverse autophagy defects in patient-derived mDA progenitors

We then utilised this high-content imaging LC3 assay to identify novel compounds of potential therapeutic interest for BPAN by screening the Prestwick Chemical Library containing 1,280 compounds, of which more than 95% FDA/ EMA approved (Fig. 4A). DMSO and Bafilomycin A1 were used as negative and positive controls, respectively. Image analysis and hit Z-score calculation was performed to rank the hits (Fig. 4B).^60^ Several compounds, that enhanced LC3 puncta number at levels higher than the positive controls, were identified (Fig. 4C-E).

**Figure 4.**
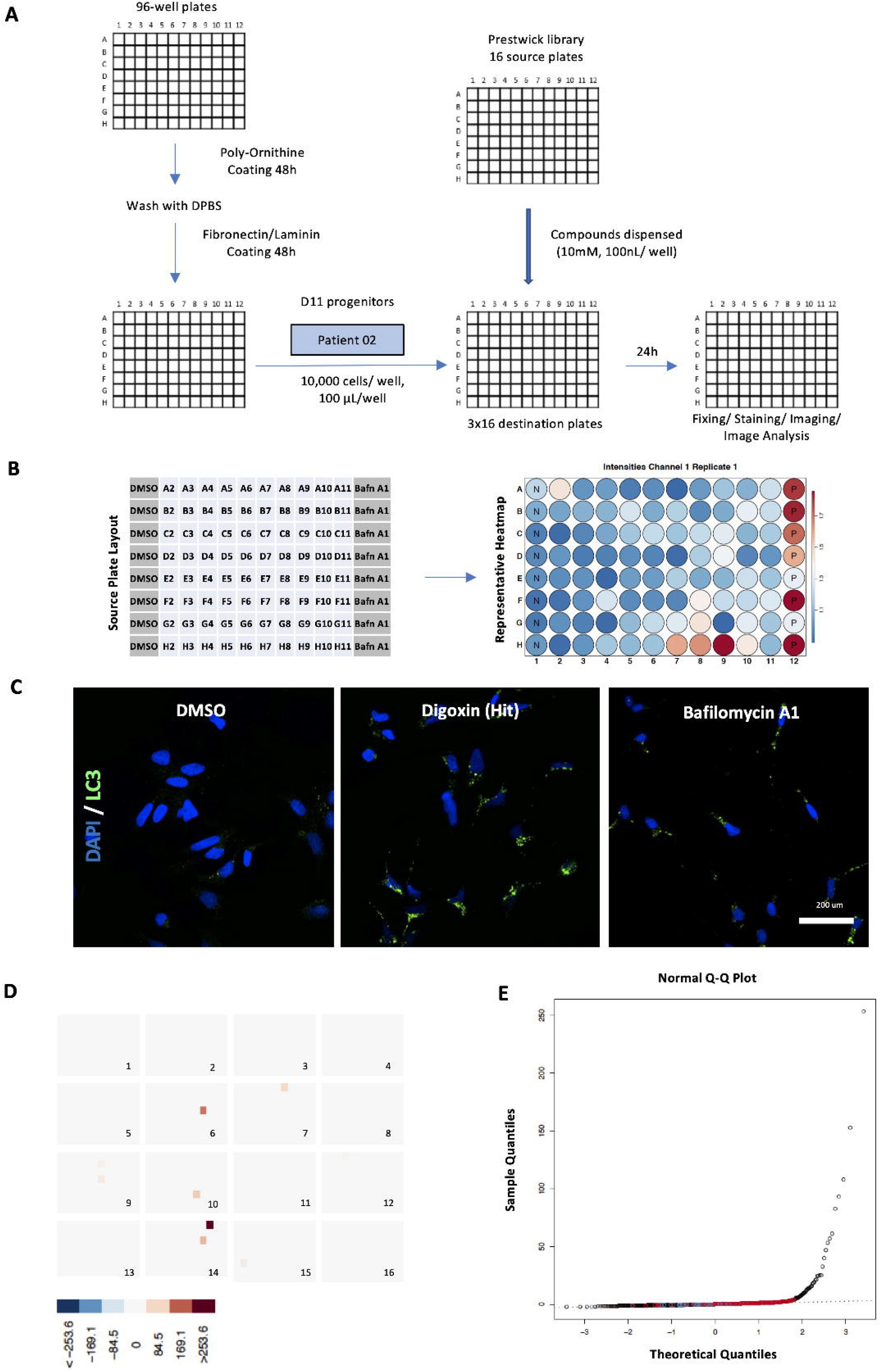
Drug Screening. **(A)** Screening protocol. Patient 02 cells used for the screen. The process was optimised and, wherever possible, automated to allow large-scale compound testing. Overall, 16 48x 96-well destination plates were required (16 source plates, 80 compounds each, tested in technical triplicates). **(B)** Source plate layout (left) and a representative heatmap (right) are shown. N and P represent negative and positive controls, respectively. Increasing puncta numbers per nuclei (in hits and positive controls) are depicted in increasingly darker shade of red. **(C)** Representative images. Immunofluorescence analysis of LC3/ DAPI after treatments with DMSO, compound hit digoxin and Bafilomycin A1. Digoxin increases LC3 puncta production to levels higher than in Bafilomycin-treated cells. Scale bar, 200 µm **(D)** Distribution of hits with the highest z-scores. There is an even distribution across all 16 source plates. **(E)** Quantile-Quantile plot for screened compounds. Several compound hits have z-scores and sample quantiles that are much higher than their theoretical values, as well as values from positive controls (in red).

The hits identified in the Prestwick library screen were further validated through a second high content imaging-based assay examining p62/SQSTM1 degradation (Fig. 5A, 5B). P62/SQSTM1 is a cargo receptor that attracts the machinery to protein aggregates and can also been used as a marker of autophagy flux. Combined degradation of p62/SQSTM1 and LC3 accumulation point towards autophagy flux induction, whereas increases in both puncta numbers indicate a flux block.^57, 58, 61^ For the p62/SQSTM1 validation assay, hits leading to LC3 puncta increase with the 26 highest Z-scores were tested (Supplementary Table 9). DMSO and Torin 1 were used as negative and positive controls, respectively. Five compounds with true autophagy flux-inducing properties were identified, leading to both LC3 puncta accumulation and p62/SQSTM1 degradation, with no adverse effect in cell viability (Fig. 5C). All five compounds were members of the class of cardiac glycosides. The other compounds were either autophagy blockers or resulted in poor cell viability, and hence excluded from further validation (Fig. 5C).

**Figure 5.**
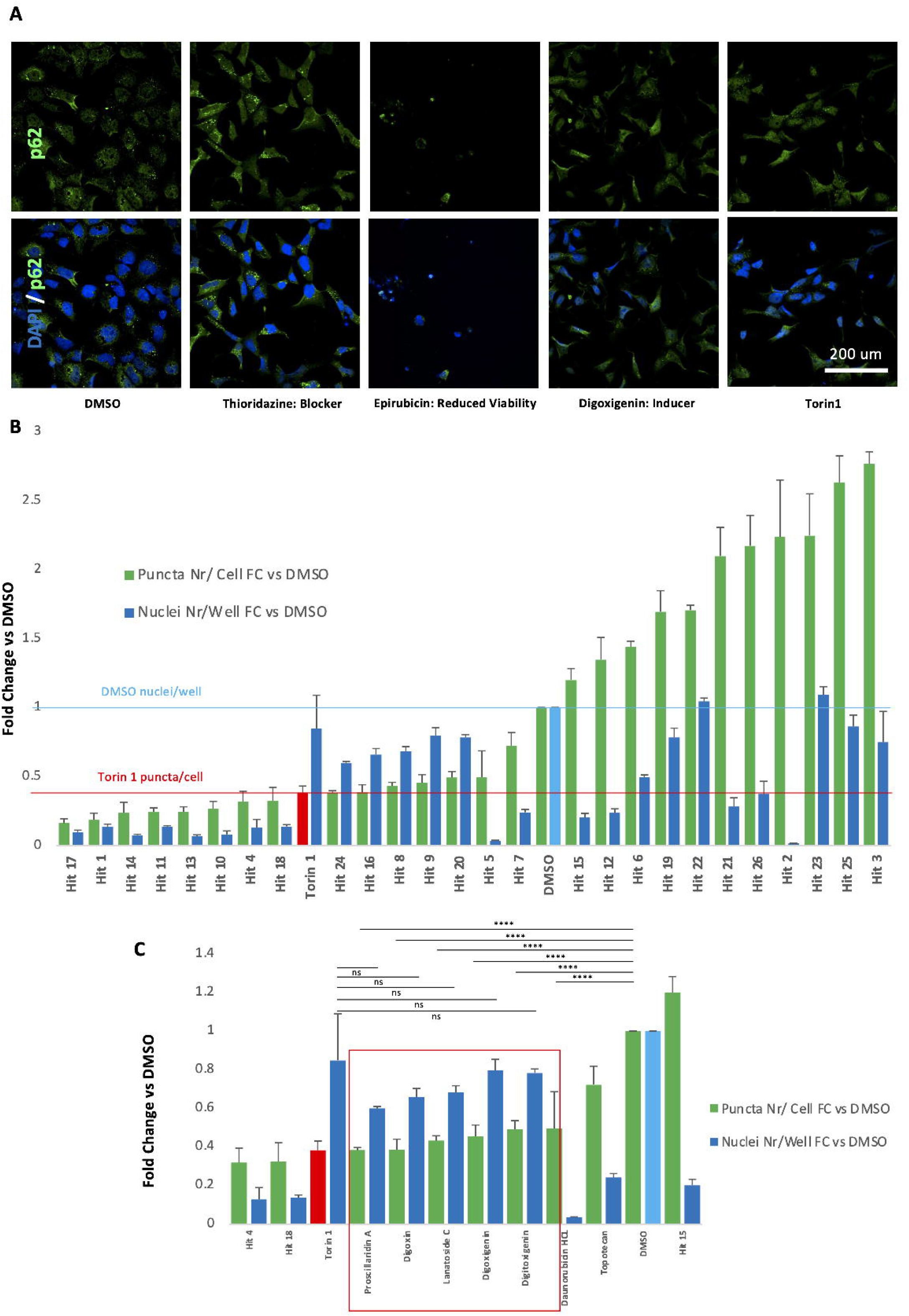
P62 hit validation assay. **(A)** Immunofluorescence analysis of p62/DAPI in Patient 02 Day 11 ventral midbrain progenitors after treatments with Prestwick screen hits with the 26 highest z-scores. Representative images from negative controls (DMSO), positive controls (Torin 1), autophagy inducers and blockers, as well as ‘false hits’ (resulting in reduced cell viability). **(B)** Quantification of cell viability (fold change of nuclei number per well, compared to DMSO) and p62 degradation (fold change of p62 puncta per nuclei, compared to DMSO; green bars) for each hit, as well as Torin 1 and DMSO. **(C)** 5 cardiac glycosides significantly degrade p62 when compared to DMSO and are associated with acceptable cell viability. There is no statistical difference in p62 puncta per cell, or nuclei per well, when comparing Torin 1 and the 5 cardiac glycosides. There is also no statistical significance between DMSO-treated and glycoside-treated nuclei per well (statistics not shown). n=3 technical replicates for each compound, n=9 for DMSO and Torin 1. Error bars represent Standard Deviation. Statistics were calculated using ANOVA. Abbreviations: FC = fold change, Nr= number

### Cardiac glycosides correct disease-specific gene expression profiles affecting key intracellular pathways in BPAN mDA neurons

Therapeutic efficacy of the identified cardiac glycosides was further tested by examining their effects on gene expression in Day 65 mDA neurons (Fig. 6, Supplementary Fig. 3E-F). Patient 02 mDA cultures were treated for 48 hours with cell culture media containing either Torin 1 or digoxin (at 250 nM and 10 μM, respectively), before harvesting at day 65 of differentiation. Bulk RNAseq analysis of the compound-treated Patient 02 cultures revealed 584 (428 overexpressed and 156 underexpressed) and 226 (84 overexpressed and 142 underexpressed) genes after treatments with digoxin and Torin 1, respectively (Fig. 6A). Comparison of gene expression profiles in Patient 02, CRISPR01 (Patient 02 Corrected) and compound-treated mDA neurons (Fig. 6B) identified correction of gene expression profiles in several previously defective intracellular pathways, including cellular response to amino acid stimuli (GO:0071230, p=1.28×10^−4^; *COL1A1, COL3A1, COL1A2, COL4A1, MMP2, COL5A2*), apoptosis (GO:0043066, *p*=0.02; *PDGFRB, ANXA1, PLK2, PDE3A, TAX1BP1, VEGFB, SPRY2, RPS27A, THBS1, CD44, MCL1*), Parkinson’s Disease (KEGG:hsa05012, *p*=0.01; *NDUFB9, NDUFA4, NDUFA3, NDUFC2, UQCRC2, COX5B, SLC25A5*) Huntington’s Disease (KEGG:hsa05016, *p*=0.01; *NDUFB9, NDUFA4, NDUFA3, NDUFC2, UQCRC2, COX5B, SLC25A5, BBC3*), mitochondrial function (GO:0006120, *p*=0.01; *NDUFB9, NDUFA4, NDUFA3, NDUFC2*), calcium signalling (GO:0051592, *p*=0.02; *PENK, ADAM9, ANXA7, THBS1*) and collagen-related processes (GO:0030199, *p*=5.79×10^−9^; *COL1A1, COL3A1, TGFB2, COL1A2, COL5A1, LOX, LUM, COL14A1, COL5A2*). Moreover, examining the specific effect of digoxin treatment on BPAN mDA neurons revealed overexpression of several mitochondrially-encoded genes involved in the mitochondrial electron transport chain^62^, such as *MT-CO1, MT-CO2, MT-CO2, MT-ND4, MT-ND5, MT-ND6* (Fig. 6A).

**Figure 6.**
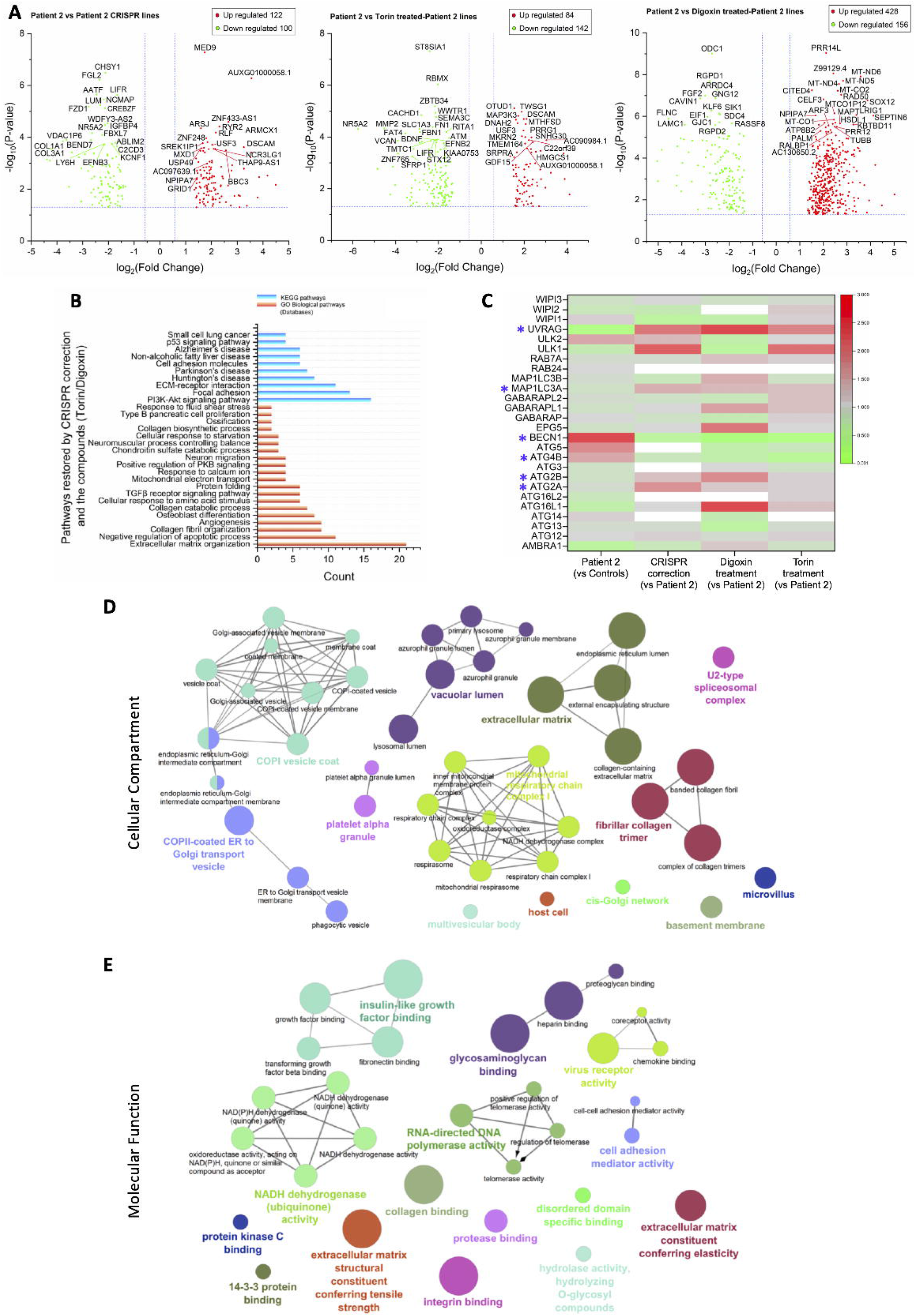
Correction of intracellular pathways after compound hit 48-hour treatment. **(A)** Volcano plots comparing Patient 02, Patient 02 compound-treated and Patient 02 CRISPR-corrected (CRISPR 01) mDA neuronal gene expression. **(B)** Intracellular pathways jointly corrected by both *WDR45* CRISPR-mediated mutation correction and compound treatments. **(C)** ATG differential expression and correction after CRISPR-genome editing and compound treatments. ATGs with consistent differential (over- or under) expression after CRISPR correction and compound treatments, when compared to untreated Patient 02 neurons, are marked with *. White= not mapped during RNAseq. **(D), (E)** ClueGO analysis of GO term enrichment of differentially expressed genes, showing network graphs of differentially expressed genes between Patient 02, CRISPR01, and compound-treated Patient 02 mDA neurons. Cellular components (CC) (C), and molecular functions (MF) (D) are shown. Network graph nodes represent GO terms (the most significant are named) and edges indicate shared genes between GO terms. Functional groups of GO terms are indicated by the same colour. GO functional groups exhibiting statistically significant differences (p< 0.05) are shown. n=3 biological for all lines, analysis based on median TPM values. *P*-values of <0.05 and fold change of <0.5 or >2 [-1<log2(FC)>1] (Student’s *t*-test) were set as statistically significant cut-offs. The top 40 genes (as per lowest *p*-values) are labelled in volcano plots.

Expression of autophagy-related genes (ATG) was also restored after compound treatments. Bulk RNA sequencing data were re-interrogated, without applying *P*-value and fold change cut-offs. The effect of digoxin and Torin 1 in ATG expression in Patient 02 Day 65 mDA neurons (Fig. 6C, Supplementary Table 10) was examined; ATG expression profiles in the corresponding isogenic control (CRISPR 01) were also compared. Firstly, WDR45 was over-expressed in Patient 02 compared to Controls 01 and 02, but unmapped in the other conditions. Moreover, *BECN1* (encoding Beclin 1, which forms a complex with vps34/PI3K class III and is essential for PI3P production and autophagosome formation)^8^ was overexpressed in untreated Patient 02 mDA neurons (fold change 4.09 versus Controls) and underexpressed in CRISPR 01 and digoxin- and Torin 1-treated lines (fold changes 0.44, 0.09 and 0.15 versus untreated patient neurons, respectively). *UVRAG* (encoding a protein that binds to the Beclin 1/ PI3K class II complex and induces autophagy)^63^ was underexpressed in untreated Patient 02 mDA neurons (fold change 0.17 versus Controls) and overexpressed in CRISPR 01 and digoxin- and Torin 1-treated lines (fold changes 2.36, 3.92 and 2.13 versus untreated patient neurons, respectively). *MAP1LC3A* (encoding an LC3 isoform), *ATG2A* and *ATG2B* were also downregulated in untreated patient neurons, but upregulated in other conditions. Conversely, *ATG4B,* important in LC3 processing,^64^ exhibited the opposite pattern, with overexpression in the untreated Patient 02 line and underexpression in CRISPR-corrected and compound-treated cells. Differentially expressed genes after compound treatments were associated with several cellular compartments, most notably coat protein complex I (COPI) and ER-Golgi transport vesicles, the vacuole lumen of lysosomes and azurophil granules, multivesicular bodies, mitochondria, the extracellular matrix and fibrillar collagen trimers, as well as the spliceosome (Fig. 6D). Restored molecular functions after compound treatments included NADH dehydrogenase activity, molecule binding, and extracellular matrix tensile strength and elasticity (Fig. 6E).

## Discussion

BPAN is a devastating childhood-onset neurodegenerative condition characterised by brain iron accumulation, for which there are currently no disease-modifying therapies^2, 3, 4, 5^. We have developed a mDA model of disease which has not only provided further insight into disease pathogenesis but also potentially identified future candidate compounds for therapeutic development. We utilised an iPSC-derived mDA model for several reasons: firstly, BPAN predominantly affects the brain with particular involvement of the substantia nigra and clinical evolution of parkinsonism in late stage disease; secondly, the mDA progenitor platform is easily amenable to screening technologies such as the high content imaging assay; and thirdly, it facilitates preclinical therapeutic testing in a humanised model of disease, relating to specific patient mutations and genetic background. We thus envisage that use of this model to identify promising drug candidates has direct translational potential.

Several measures were taken to overcome the recognised limitations of iPSC-based disease modelling. The inherent issue of iPSC line variability^65^ was countered by using multiple patient and control lines, including CRISPR-corrected isogenic controls. Moreover, the XCI status of female lines was confirmed ensuring that only the mutant *WDR45* allele was expressed in all differentiated patient lines. Stringent characterisation of all iPSC lines and mDA cultures also ensured reproducible results in downstream experiments. As a result, the model recapitulated key disease-related features including defective LC3 processing and aberrant gene expression profiles in several disease-related intracellular pathways, such as neurodevelopment and ferroptosis; the latter has been proposed as a mechanism leading to disease in BPAN.^66^ Examination of specific differentially expressed genes also revealed interesting links to BPAN pathophysiology. For example, underexpression of *HCRT* in BPAN lines fits with the clinical observation that BPAN patients often present with disordered sleep and hyperphagia.^52^ Moreover, several dysregulated genes have been linked with intellectual disability (*PCDHGA10*),^43^ autism (*FAM120C*)^44^ and aberrant CNS development (*MAPK7, PCDHGA10, PTPRT, LRFN1*),^45, 46, 53, 54^ all of which feature prominently in BPAN. Notably, collagen metabolism pathways were also prominently involved in our BPAN disease model, with interesting links to neurodegeneration.^67, 68, 69, 70^ Lack of COL1A1 and COL6A1 are associated with increased spontaneous apoptosis, abnormal autophagy and increased vulnerability to oxidative stress.^67, 70^ Genome-wide association studies have identified a link between dementia and abnormal collagen I metabolism. Accumulation of alpha-synuclein and ER stress through COL4A2 dysregulation is also reported.^68^ Many collagen-related genes were specifically underexpressed in BPAN mDA neurons with restoration of expression after CRISPR correction and compound treatments. Our data therefore supports a role for collagen metabolism in BPAN pathogenesis, which warrants further investigation.

Overall, it is not clear how WDR45 deficiency leads to dysregulation of gene expression. Our RNAseq dataset implicates a wide network of WDR45-interacting proteins in adjacent or downstream pathways, or may also point towards a role of WDR45 in regulating gene expression. Irrespective of the underlying mechanism, GO Term and KEGG pathway analysis identified disturbances in key intracellular pathways implicated in BPAN by other studies (REF).^21, 22, 66^ More functional analysis at the protein level is warranted in future studies to elucidate the exact intracellular role of WDR45 in neurodevelopment and neurodegeneration.

LC3 puncta formation was also defective both in BPAN fibroblasts and mDA progenitors, both at basal conditions and after autophagy flux induction or inhibition. However, more work is warranted to elucidate the exact molecular mechanisms underpinning this flux defect and understand how lead compounds may reverse it. There was also evidence of abnormal ATG expression in BPAN mDA neurons. Several genes involved in the early stages of autophagosome formation (e.g., *BECN1, UVRAG, ATG2A/B, ATG4B, MAP1LC3A*) were differentially expressed when compared to controls. The expression pattern was restored after CRISPR genome editing and, importantly, also after compound treatments. Some genes were not mapped during the RNAseq and further studies examining ATGs at the protein level are warranted; nevertheless, our findings suggest a dysregulated ATG network in BPAN neurons with potential for correction after targeted therapeutic interventions.

The pathophysiology of BPAN has been studied in other cellular^5, 7, 22, 71^- and animal^6, 18, 21^ models and our work corroborates and builds upon many of these findings. In mice, WDR45 deficiency leads to defects in autophagy-mediated clearance of protein aggregates^6^ and accumulation of neuronal ER stress, with subsequent apoptosis.^21^ Seibler and colleagues^22^ report patient fibroblasts and iPSC-derived mDA neurons with reduced membrane-bound LC3 (LC3-II) - pointing towards defective autophagy - as well as iron accumulation and possible mitochondrial and lysosomal dysfunction. Iron accumulation may be in part, attributed to impaired autophagic degradation of the transferrin receptor, which in turn might lead to ferroptosis.^66^ There is some indication that autophagy induction in BPAN cells might lead to reversal of iron accumulation and ER stress.^21, 22^ Our study corroborates pertinent WDR45-specific findings in the Seibler study, also using more patient lines and two age-matched controls; moreover, we’ve been able to better understand gene expression associated with WDR45 deficiency and develop potential novel autophagy-enhancing therapies. Overall, both our own data and published work indicate that autophagy is likely to be a key pathway for therapeutic targeting in BPAN.

Whilst iPSC-based models are increasingly used for modelling neurological disorders, their use as a platform for drug screening and novel therapeutic development is still relatively rare.^72, 73, 74, 75^ This may be in part attributed to a number of technical hurdles, related not so much to scaling up production of iPSCs and differentiated cells, but to designing large-scale assays, generation of big datasets, and automation.^74^ In our study, we conducted the Prestwick screen using one patient line, and testing each drug in triplicate destination plates required 48 x 96-well destination plates. To address this volume, the screening protocol (cell seeding, fixing, staining and subsequent image acquisition and analysis) was automated, wherever possible.^75^ The FDA-approved Prestwick library was screened to identify compounds for potential repurposing in BPAN. Cardiac glycosides were identified as the top candidates for future therapeutic development, acting as potent autophagy inducers in BPAN patient cells with no obvious adverse effects in cell viability. Furthermore, treatment of patient mDA neurons with Torin 1 and digoxin also normalised gene expression in a number of genes involved in defective pathways related to autophagy, Parkinson’s and Huntington’s disease, apoptosis, mitochondrial function, calcium signalling and collagen-related processes. These pathways have been implicated in neurodegenerative processes,^76, 77, 78^ hence this restoration of gene expression, as well as the digoxin-mediated overexpression of mitochondrially-encoded genes involved in the electron transport chain, provides further validation of efficacy for the tested compounds.

Cardiac glycosides are of particular interest from a mechanistic perspective. Firstly, although they have been identified as potential activators of autophagy,^79, 80, 81^ their exact role in regulating this process is not well understood. Cardiac glycosides inhibit the Na(+)/K(+)-ATPase and may induce autophagy by activating AMPK (and inhibiting mTOR) or by regulating ERK1/2 signalling.^82^ Interestingly, WDR45 forms a complex with ATG2A which also contains AMPK and ULK1 in normal conditions. Upon starvation - possibly due to AMPK activation - the WDR45/ ATG2A complex dissociates from AMPK and ULK1 and localises to the forming autophagosome and promotes its maturation^7^. Other studies have suggested a different mechanism of action, namely that cardiac glycosides may reduce autophagy flux, and thus prevent autophagy-dependent cellular death under conditions of starvation or hypoxia and ischaemia.^81^ In our work, cardiac glycosides had autophagy-inducing properties but further studies, especially application of stressors, might be warranted. Regardless, studies have highlighted the neuroprotective properties of cardiac glycosides in hypoxia and glucose starvation,^83^ and indeed the latter is thought to be a key stimulus for intracellular WDR45 activation.^7^ Cardiac glycosides are widely used licensed drugs with established pharmacokinetics and side effect profiles; as such, they could be considered for future drug repurposing trials in BPAN. The potential bottleneck of blood brain barrier impenetrability^83, 84^ could be overcome by the use of structurally related compounds with better blood brain barrier permeability (e.g. digitoxin, neriifolin),^84, 85^ compound modification, or even intrathecal/intracerebral drug delivery.

In conclusion, we describe the successful generation of a humanised iPSC-based mDA neuronal model of BPAN, which provides important insights into the dysregulated cellular pathways and disease pathophysiology underpinning this condition. Moreover, the use of this model as a platform for a high content imaging-based drug screen has identified compounds that have clear potential for development as new, targeted and efficacious treatments.

## Materials and Methods

### Patient ascertainment

Written informed consent was obtained from all participants for the use of fibroblasts for this study. Two age-matched healthy control fibroblast lines were ascertained from the Dubowitz biobank (MTA/IDT # 38).

### IPSC generation and maintenance

iPSC lines were generated^23^ from patient and control dermal fibroblasts isolated from skin biopsies, as approved by the Local Research Ethics Committee (Reference 13/LO/0171) and UCL GOS Institue of Child Health (R&D Reference 14NM20)(Supplementary Table 1).

Fibroblasts were maintained in DMEM (Gibco), 10% foetal bovine serum, (FBS, Gibco), 2mM L-glutamine (Gibco), 1% MEM non-essential amino acids (Gibco), and 1% penicillin/streptomycin (Gibco). Reprogramming into iPSC was performed using the CytoTuneVR -iPS 2.0 Sendai Reprogramming kit (Invitrogen).^26, 86^ Derived iPSCs were maintained in mTeSR1 (StemCell Technologies) on Matrigel (Corning)-coated plates, and regularly passaged with EDTA, 0.02% solution (Sigma-Aldrich). Two isogenic control lines were created by CRISPR-Cas9 correction of the c.19C>T and c.700C>T variants in patients 2 and 3, respectively (Supplementary Table 1). iPSC lines from each patient, age-matched healthy controls and CRISPR-corrected lines, were characterised for pluripotency. Afterwards, one clone per patient and control were used for downstream experiments.

### Generation of CRISPR/Cas9-corrected iPSC lines

Single guide RNAs (sgRNAs) were designed to target the nonsense variants of interest in the *WDR45* gene, using Welcome Sanger Institute Genome editing website (https://wge.stemcell.sanger.ac.uk/) (Synthego) (Supplementary Table 2).

For the Homology-Directed Repair (HDR) donor template (Integrated DNA Technologies, Inc), 90bp long single-stranded oligodeoxynucleotides (ssODNs) were designed, containing the missense variant and 2 silent mutations (to avoid repeated editing once the desired correction had occurred) (Supplementary Table 2). Prior to nucleofection, IPSC culture medium was supplemented with 10ug/ml ROCK Inhibitor Y-27632 (Selleckchem) for 24hrs. 20 μg recombinant *Streptococcus pyogenes* Cas9 protein (department of Biochemistry, university of Cambridge) was combined with 225 μΜ full length sgRNA (Synthego) and incubated at room temperature for 10 minutes. 500 μM of HDR donor template was added to the preassembled Cas9 ribonucleoprotein. Cells were dissociated with StemPro™ Accutase™ (Gibco), and 1 x 10^6^ cells were combined with the preassembled Cas9 ribonucleoprotein with the donor DNA mix, nucleofected in a 100 µl reaction using the Lonza 4D-Nucleofector System with P3 Primary Cell 4D-Nucleofector X Kit (Lonza) and program CA137. Nucleofected cells were plated onto 10cm dishes in medium containing 10 uM Y-27632 and incubated at 37 °C under 5% CO2. Edited cells were expanded to approximately 70% confluency before subcloning. Approximately 300 cells were subcloned onto 10 cm dishes. After 14 days, clonal colonies were picked, expanded and genotyped.

### iPSC mDA neuronal differentiation

iPSCs were differentiated into mDA neurons as previously described.^26, 27, 86, 87^ iPSCs were harvested using TrypLE^TM^ (ThermoFisher) and plated onto non-adherent bacterial dishes (Corning) (1.5×10^5^ per cm^2^), in DMEM/F12:Neurobasal (1:1) (ThermoFisher), N2 (1:100, Thermo Fisher), B27 minus vitamin A (1:50) (Invitrogen), 2 mM L-glutamine (ThermoFisher) and ROCK-inhibitor (Thiazovivin, Cambridge Bioscience) for the first 2 days. Purmorphamine (0.5 μM, Cambridge Bioscience) was added to the medium from day 2 to day 9 and Thiazovivin was removed on day 2. On day 4, EBs were plated onto pre-coated with polyornithine (PO; 15 μg/ml; Sigma), fibronectin (FN; 5 μg/ml Gibco) and laminin (LN; 5 μg/ml; Sigma) 12-well plates (Corning) in DMEM/F12:Neurobasal (1:1), N2 (1:200), B27 minus vitamin A (1:100), 2 mM L-glutamine. From day 0 to day 9 medium was supplemented with 100 nM LDN193189 (Stemgent Inc.), 0.8 µM CHIR99021 (Tocris Bioscience) and 100 ng/ml hSHH-C24-II (R&D Systems) and also with 10 μM SB431542 (Tocris Bioscience) from day 0 to day 6. On Day 11, cells were re-plated on PO/FN/LN 12-well plates in droplets of 1.5 ×10^4^ cells per µl, in medium including Neurobasal B27 minus vitamin A (1:50), 2 mM L-glutamine, 0.2 mM ascorbic acid (AA) (Sigma-Aldrich) and 20 ng/ml BDNF (Miltenyi Biotech). From day 14 to day 65, medium was supplemented with 0.5 mM dibutyryl c-AMP (Sigma-Aldrich) and 20 ng/ml GDNF (Miltenyi Biotech). On day 30 of differentiation, cells were re-plated as described for day 11 and γ-secretase inhibitor DAPT (10 μM, Tocris) was added to the medium. Cells were harvested at either day 11 or 65 for analysis.

### DNA and cDNA extraction

DNA was extracted using a commercial kit (DNeasy Blood & Tissue kit, Qiagen). For cDNA, total RNA extraction from pellets was performed using the RNeasy Mini Kit (Qiagen). RNA was purified using the DNase I Kit (ThermoFisher), while reverse transcription took place via the Superscript III Reverse Transcriptase kit (ThermoFisher).

### *WDR45* genomic and cDNA sequencing

Direct Sanger Sequencing of genomic DNA extracted from patient-derived iPSC lines and CRISPR-corrected lines was performed to confirm genotype. cDNA was also sequenced to determine which of the two *WDR45* alleles, the mutated or wild-type, was expressed in the two female iPSC lines (Patient 01 and Patient 03).

DNA sequences were obtained from the Genome Reference Consortium Human Build 37 (GRCh37), (https://grch37.ensembl.org/Homo_sapiens/Info/Index) (transcript ID ENST00000356463.3, NM_007075). Primers were designed with Primer3Plus software and genomic DNA/ cDNA was amplified by PCR (Supplementary Tables 3 and 4). Amplified DNA/ cDNA was then purified with MicroCLEAN Kit (Clent Life Science) and further processed with the BigDye Terminator v1.1 Cycle Sequencing Kit (ThermoFisher). Sequencing was performed with the ABI PRISM 3730 DNA Analyzer (Applied Biosystems). Results were analysed using MutationSurveyor^®^ (https://www.softgenetics.com/mutationSurveyor.php). For sequence alignment, the wild type *WDR45* sequence was obtained from the Ensembl genome browser, amino acid sequence predictions were generated by entering wild type, patient and CRISPR-corrected sequences into the online ExPASy Translate tool (https://web.expasy.org/translate/), and sequence alignments were generated using the online ClustalOmega tool (https://www.ebi.ac.uk/Tools/msa/clustalo/).

### X Chromosome Inactivation (XCI) Studies

Genomic DNA from fibroblasts, iPSCs and mDA neurons deriving from female BPAN patients was extracted using the DNeasy Blood & Tissue Kit (Qiagen). XCI status was studied by examining differences in methylation at the human Androgen Receptor (AR) locus located on the X Chromosome (North East Thames Regional Genetics Laboratory at Great Ormond Street Hospital). The assay is based on the activity of the methylation-sensitive restriction enzyme HpaII, allowing subsequent PCR amplification and detection of a CAG repeat-containing portion of the methylated (inactivated) AR allele.^32^

### Assessment of genome integrity

Genome integrity was assessed by the Infinium HumanCytoSNP-12 v2.1 BeadChip array (Illumina) using genomic DNA from iPSC lines (UCL Genomics). The BlueFuse Multi software (https://www.illumina.com/clinical/clinical_informatics/bluefuse.html) (Illumina) was used to generate karyograms. All clones with large (> 5 Mb) and/ or pathogenic deletions/ gains were excluded from downstream experiments.

### Epi-Pluri-Score

Genomic DNA from iPSC lines was additionally analysed with Epi-Pluri-Score (Cygenia), which compares pluripotent with non-pluripotent cells. DNA-methylation levels (β-values) were measured by pyrosequencing at three specific CpG sites, within the genes *ANKRD46*, *C14orf115*, and *POU5F1 (OCT4)*. The CpG sites at *ANKRD46* and *C14orf115* in pluripotent cells are methylated and non-methylated, respectively, and these β-values were combined into the Epi-Pluri-Score. Methylation within *POU5F1* might indicate early differentiation, so this β-value was also factored in.^25^

### *In vitro* spontaneous differentiation

To assess iPSC pluripotency, embryoid bodies (EB) were generated by harvesting iPSCs with TrypLE and plating them onto non-adherent bacterial dishes at a concentration of 1.5 x 10^5^ per cm^2^, in Knockout-DMEM, 20% Knockout-Serum Replacement (Invitrogen), L-glutamine (2 mM; ThermoFisher), 1% Non-Essential Amino Acids 100x (Invitrogen), 50 μM 2-Mercaptoethanol (ThermoFisher), plus 1 μM Thiazovivin for the first 2 days. On day 4, EBs were seeded into a 24-well plate (Corning) for spontaneous differentiation into all three germ layers. For neuroectoderm and endoderm differentiation, wells were coated with matrigel and cells maintained in the same medium, as above. For mesodermal differentiation, wells were coated with 0.1% gelatin (Sigma-Aldrich) and cells maintained in DMEM supplemented with 20% FBS and 2 mM L-Glutamine.

On Day 16, cells were analysed by immunofluorescence for expression of endoderm-, mesoderm- and ectoderm-related markers.

### Reverse Transcription (RT) PCR

Detection of the Sendai virus (SeV) genome and SeV-carried pluripotency genes in iPSCs occurred through PCR using cDNA from each iPSC line and primers provided by the CytoTune-iPS 2.0 Sendai Reprogramming Kit (Invitrogen). PCR was also performed on cDNA from each iPSC line in order to detect expression of pluripotency markers *SOX2, cMYC, NANOG, OCT4 and KLF4* (Supplementary Tables 5 and 6).

### Real-time quantitative RT PCR (qRT PCR)

qRT-PCR reactions were prepared using 1x MESA Blue qPCR MasterMix Plus for SYBR® Assay (Eurogentec), 0.1 µl ROX Reference Dye (Invitrogen), 9 μL cDNA (dilution 1:25) and 500 nM of each primer. qRT-PCR was performed in technical triplicates with the StepOnePlus Real-Time PCR System (Applied Biosystems) (Supplementary Table 7). Relative gene expression to the housekeeping gene glyceraldehyde-3-phosphate dehydrogenase (*GAPDH*) was analysed using the 2^-ΔΔCt^ method.

### Immunocytochemistry

Cells were washed in 1x DPBS, fixed with 4% Paraformaldehyde (PFA) (Santa Cruz), then blocked [in 1x DPBS, 10% FBS, with or without 0.1% of 100X Triton for cell membrane permeabilisation (for nuclear/cytoplasmic and membrane markers, respectively; for Day 65 mDA neurons only, 0.3% Triton was used], incubated with primary antibody solution (in blocking buffer, at +4°C overnight), washed again and, finally, incubated with the secondary antibody solution (and DAPI 1 mg/ml) at room temperature for 1 hour.

Imaging was performed with the Olympus IX71 inverted TC scope for assessment of iPSCs (pluripotency marker expression, spontaneous *in vitro* differentiation) and day 11 mDA precursors. A multiphoton confocal microscope (Zeiss LSM880) was used for Day 65 studies.

For quantification, 4 random fields were imaged from each independent experiment. Subsequently, 1200 to 1800 randomly selected nuclei were quantified using ImageJ (National Institutes of Health). Manual counting for nuclear (DAPI) staining and co-staining with the marker of interest was performed, and percentages of cells expressing combinations of markers were calculated as needed.

### High content imaging-based immunofluorescence

Day 11 mDA progenitors were plated onto 96-well low skirted, PO/FN/LN-coated, microplates (CellCarrier-96 Ultra Microplates, PerkinElmer) at a density of 15,000 cells/ well. For each biological replicate, all 7 lines had the same start date of differentiation and were seeded on the same 96-well plate. Cells were treated 24 hours post-plating, then fixed with 4% PFA, washed, permeabilised and fixed a second time in ice-cold methanol for 15 minutes at +4°C, quenched in 50 mM NH_4_Cl in DPBS for 20 minutes, blocked (in DPBS, 10% FBS and 0.1% Triton) for 1 hour at room temperature and, finally, incubated with primary and secondary antibodies as stated above.

Treatments for the LC3 assay included 3 hours with mTOR inhibitor Torin1 (Merck-Millipore, final concentration 250 nM), Bafilomycin A1 (Sigma, final concentration 10 nM) or dimethyl sulfoxide (DMSO, Sigma-Aldrich; 1:1000 dilution). All treatments were performed in technical duplicates.

For drug screening, the Prestwick Chemical Library (1,280 compounds, 95% FDA/ EMA approved, 10 mM in DMSO, https://www.prestwickchemical.com/screening-libraries/prestwick-chemical-library/) was used; cells were treated with compounds for 24 hours at 10 μM final concentration.

Patient 02 (587-02) cells were used and 48x 96-well destination plates were required (16 source plates, 80 compounds each, tested in triplicates). Plates were automatically coated and washed using the Beckman Coulter Biomek NXp and Biotek EL406 workstation, automated by the SAMI scheduling software (Beckman Coulter). Cell seeding onto 96-well plates (and the subsequent PFA fixing, post drug treatments) was performed with the Biotek Multi-Flo Multi-Mode Dispenser. For drug dispensing from source into destination plates, the Labcyte Echo 550 with CellAccess system was used. For immunostaining, the combination of the Beckman Coulter Biomek station and the SAMI scheduling software was utilised again.

For all high content imaging-based experiments, the PerkinElmer Opera Phenix microscope was used for imaging. 20 fields were imaged per well, at 40 x magnification, Numerical Aperture 1.1, Binning 1. Image analysis was performed using ImageJ and R Studio.^60^ For the drug screen, puncta values were normalised according to positive and negative controls from each plate and Z-scores for each compound screened were generated. Statistical significances were calculated on GraphPad Prism V. 8.1.2. software (GraphPad Software, Inc.; https://www.graphpad.com/scientific-software/prism/).

### Immunoblotting

Proteins were extracted from cell pellets resuspended in RIPA buffer (Sigma) containing cOmplete^TM^ Mini Protease Inhibitor Cocktail (Merck) (1:10 dilution). Protein concentration was quantified using the Pierce BCA Protein Assay Kit (ThermoFisher). 10 μg of protein were mixed with 4x Laemmli Buffer (Bio-Rad Laboratories LTD), 0.1 M Dithiothreitol (DTT) (Sigma), and H20 added for equal volume loading. Samples were loaded onto 4-20% Mini-PROTEAN TGX Precast Gels (BioRad) or in 1x Tris-glycine-SDS running buffer (BioRad). Proteins were transferred onto Trans-Blot® Turbo™ Mini PVDF Transfer membranes with the Trans-Blot Turbo Transfer System (Bio-Rad). Membranes were blocked with 5% skimmed milk in PBS with added Tween20 0.05% (Merck) (PBS-T) at 4**°**C overnight. The membrane was probed with primary antibody in blocking buffer at at 4**°**C overnight, washed with PBS-T, incubated with corresponding HRP-conjugated secondary antibodies in blocking buffer for 1 hour at room temperature, then washed again, developed using HRP substrate (Clarity Western ECL Substrate, BioRad), exposed and imaged using the BioRAD Gel Doc Imager and ImageLab software (BioRAD). Band intensities on exported .tiff images were quantified with ImageJ.

### Antibodies

Primary and secondary antibodies used for immunofluorescence and immunoblotting are detailed in Supplementary Table 8.

### Bulk RNAseq

Bulk RNAseq was performed on total RNA from Day 65 neurons (UCL Genomics). The mRNA library was prepared using the Kapa mRNA HyperPrep Kit (Kapa Biosystems) and sequenced on the Illumina HiSeq 3000 system. RNAseq data quality control was performed using MultiQC (SciLife, Inc.; https://multiqc.info). RNAseq raw data in FASTQ format were analysed with Subio platform v1.24.5839 (Subio, Inc., Kagoshima, Japan, https://www.subioplatform.com/) assessing differential gene expression between patients, controls, CRISPR lines, and compound-treated lines. A P-value of <0.05 and fold change of <0.5 or >2 [-1<log2(FC)>1] (Student’s *t*-test) was set as a statistically significant cut-off to generate a list of differentially expressed genes. GO enrichment analysis was performed using DAVID Bioinformatics Resources v6.8^88^ and PANTHER v16.0^89^ where a *P-*value of <0.05 was considered as significant to select enriched terms. The ontology terms biological process (BP), cellular component (CC), and molecular function (MF) were visualised in bar graphs and pie charts. ClueGo v2.5.8,^90^ a Cytoscape plugin (National Institute of General Medical Sciences) was used for the construction of the network graphs and clustering analysis.

### Quantification and statistical analysis

Statistical analysis was performed using GraphPad Prism V. 8.1.2. Patient and control samples were compared through the Student’s unpaired two tailed t-test, and the ordinary one-way ANOVA tests. On graphs, error bars represent the Standard Error of Mean. Significance levels are determined through *p-*values, and statistically significant values (*p<*0.05) are shown with asterisks, as follows; 1 asterisk (*) indicates *p-*values between 0.05 and 0.01, 2 asterisks (**) indicate *p-*values between 0.01 and 0.001, 3 asterisks (***) indicate *p*< 0.001 and 4 asterisks (****) indicate *p*< 0.0001. Nonsignificant values (*p>*0.05) are shown as (ns).

## Acknowledgements

This research was supported by the National Institute for Health Research Biomedical Research Centre at Great Ormond Street Hospital for Children NHS Foundation Trust and University College London.

The project was supported by the National Center for Advancing Translational Sciences (NCATS), National Institutes of Health, through Grant Award Number UL1TR002369. The content is solely the responsibility of the authors and does not necessarily represent the official views of the NIH.

This work was supported by a Clinical Research Training Fellowship (Action Medical Research and the British Paediatric Neurology Association, GN2465) to Apostolos Papandreou, Wellcome Intermediate Clinical Fellowship (WT098524MA to Manju A. Kurian, funding Serena Barral), Rosetrees Trust/Robert Luff Foundation (M865-CD1 to Manju A. Kurian and Apostolos Papandreou), NIHR Research Professorship (NIHR-RP-2016-07-019 to Manju A. Kurian), Sir Jules Thorn Award for Biomedical Research (to Manju A. Kurian), UK Medical Research Council funding to the MRC LMCB (MC_U12266B to Robin Ketteler), Dementia Platform UK (MR/M02492X/1 to Robin Ketteler), the Million Dollar Bike Ride Charity (University of Pennsylvania) (MDBR-19-102-BPAN, to Apostolos Papandreou and Robin Ketteler), and the NBIA Disorders Association.

We are thankful for the generation of iPSC lines by the hiPSC Core Facility at the NIHR Cambridge Biomedical Research Centre. We would like to thank Professor Sharon Tooze from the Francis Crick Institute, London, UK, for gifting us an aliquot of an anti-WDR45 rabbit monoclonal antibody which was custom-made in her laboratory. We would also like to thank Drs Tanya Singh and Daniel Little at UCL MRC Laboratory for Molecular Cell Biology for their help in RNAseq data analysis and high content imaging assay set up, respectively.

## Author contributions

Apostolos Papandreou made contributions to the conception and design of the work; to the acquisition, analysis, or interpretation of most data; and has drafted the manuscript.

Nivedita Singh performed the analysis and interpretation of the RNAseq data, drafted and revised the manuscript.

Lorita Gianfrancesco made contributions to the acquisition and analysis of data; and has revised the manuscript.

Dimitri Budinger made contributions to the acquisition and analysis of data; and has revised the manuscript.

Katy Barwick made contributions to the acquisition and analysis of data; and has revised the manuscript.

Alexander Agrotis made contributions to the acquisition and analysis of data; and has revised the manuscript.

Christin Luft made contributions to the acquisition and analysis of data; and has revised the manuscript.

Ying Shao made contributions to the acquisition and analysis of data (iPSC reprogramming and CRISPR genome editing); and has revised the manuscript.

An-Sofie Lenaerts made contributions to the acquisition and analysis of data (iPSC reprogramming and CRISPR genome editing); and has revised the manuscript.

Allison Gregory made contributions to the acquisition and analysis of data; and has revised the manuscript.

Suh Young Jeong made contributions to the acquisition and analysis of data; and has revised the manuscript.

Penelope Hogarth made contributions to the acquisition and analysis of data; and has revised the manuscript.

Susan Hayflick made contributions to the acquisition and analysis of data; and has revised the manuscript.

Serena Barral made contributions to the analysis and interpretation of data; and has substantially revised the manuscript.

Janos Kriston-VIzi performed the bioinformatic analysis and interpretation of the drug screening data, and revised the manuscript.

Paul Gissen made substantial contributions to the conception and design of the work, to the analysis, or interpretation of data, and has substantially revised the manuscript.

Robin Ketteler made substantial contributions to the conception and design of the work, to the analysis, or interpretation of data, and has substantially revised the manuscript.

Manju A. Kurian made substantial contributions to the conception and design of the work, to the analysis, or interpretation of data, and has substantially revised the manuscript.

All authors have approved the submitted version (and any substantially modified version that involves the author’s contribution to the study).

Every author has agreed both to be personally accountable for their own contributions and to ensure that questions related to the accuracy or integrity of any part of the work, even ones in which the author was not personally involved, are appropriately investigated, resolved, and the resolution documented in the literature.

---

**Supplementary figure 1.**
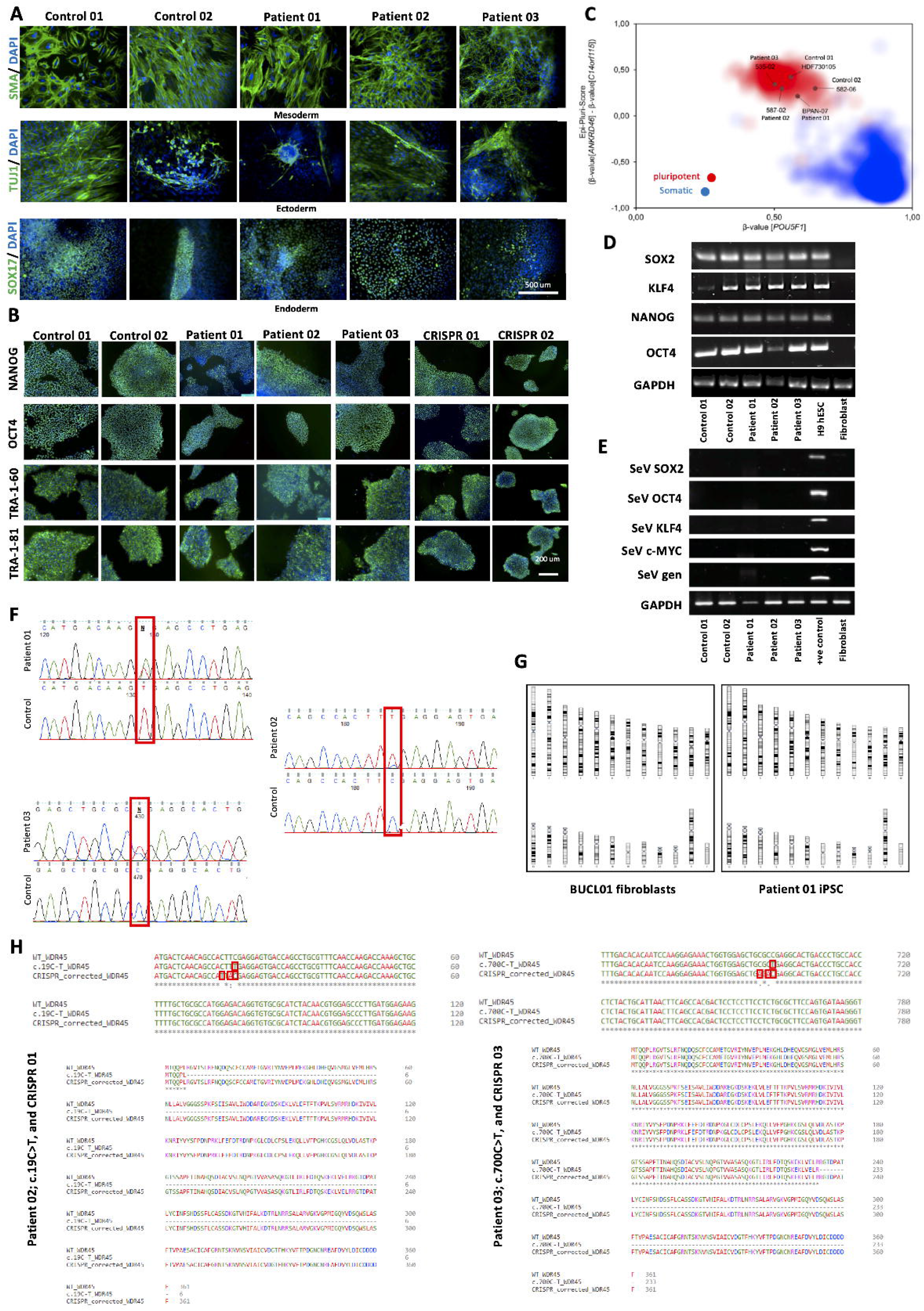
iPSC characterisation of pluripotency. **(A)** Immunofluorescence staining at Day 16 of the Spontaneous in vitro differentiation protocol. SOX17 (SRY-BOX 17; endoderm-related), TUJ1 (Neuronal Class III β-Tubulin; ectoderm-related), and SMA (Alpha Smooth Muscle Actin; mesoderm-related) markers are shown. Nuclei were counterstained with DAPI. Scale bar, 500 µm. n=1 biological replicate per line. **(B)** Immunofluorescence for pluripotency markers NANOG, OCT4, TRA-1-60, TRA-1-81 in iPSC lines. Nuclei were counterstained with DAPI. Scale bar, 200 µm. n=1 biological replicate per line. **(C)** Epi-Pluri-Score testing for iPSC lines. DNA methylation profiles (β-values) in genes ANKRD46, C14orf115, and POU5F1 for all iPSC lines match profiles of pluripotent samples (red cloud). n=1 biological replicate per line. **(D)** RT PCR in iPSC lines for expression of pluripotency markers SOX2, KLF4, NANOG, OCT4. H9 human embryonic stem cell (H9 hESC) line and human dermal fibroblasts were used as positive and negative controls, respectively. GAPDH was used as a housekeeping gene. n=1 biological replicate per line. **(E)** RT PCR in iPSC lines for detection of Sendai virus genome and pluripotency transgenes. A positive (+ve) control (SeV DNA) and a negative control (cDNA from the H9 human embryonic stem cell line, H9 hESC) were also analysed. GAPDH was used as a housekeeping gene. n=1 biological replicate per line. **(F)** Chromatograms from genomic DNA sequencing in BPAN iPSC lines. iPSC lines maintain disease-causing mutations. WDR45 disease-causing mutations are highlighted in the red rectangles. n=1 biological replicate per line. **(G)** SNP array analysis of all iPSC lines used for downstream experiments, including the two isogenic controls. Representative images. All deletions/ gains in iPSCs used for downstream experiments were small (<5Mb) and deemed as non-pathogenic by BlueFuse Multi. n=1 biological replicate per line. **(H)** Alignment of wild type, patient 02 (c.19C>T), patient 03 (c.700C>T) and CRISPR corrected WDR45 genomic DNA (above) and amino acid (below) sequences. Premature protein truncation results from both c.19C>T and c.700C>T mutations. For each CRISPR-corrected line, three nucleotide substitutions have occurred after HDR (red rectangles). For both corrections, the first two are silent/ synonymous changes and, overall, the sequence leads to translation of a full-length WDR45 protein.

**Supplementary Figure 2.**
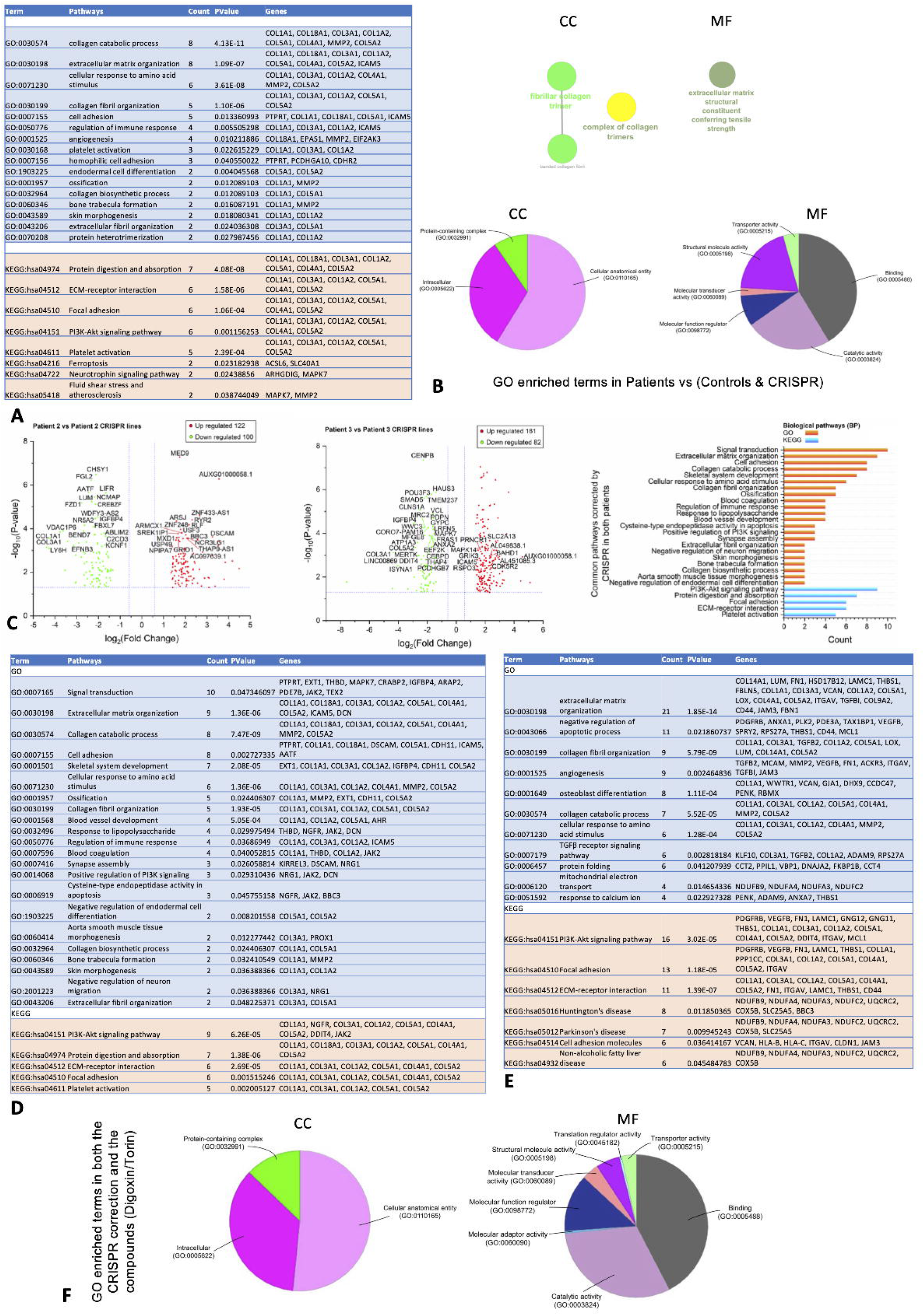
Generation and basic characterisation of mDA model. **(A)** Protocol for A9-type mDA differentiation. **(B)** Immunofluorescence for ventral midbrain progenitor-specific markers FOXA2 and LMX1A at Day 11 of mDA differentiation. Nuclei were counterstained with DAPI. Scale bar, 500 µm. n=3 biological replicates per line. **(C)** Quantification of FOXA2 and LMX1A abundance in Day 11 progenitors. n=3 biological replicates for all lines, 3 individual images from random areas of a well for each biological replicate. Percentages were calculated after manual counting of cells on ImageJ/Fiji (approximately 500 nuclei counted per image, followed by counting of cells also staining positive for FOXA2 and/or LMX1A). **(D)** qRT-PCR at d11 for pluripotency markers OCT4 and NANOG, and midbrain related markers FOXA2, LMX1A, LMX1B, EN1, EN2, relative to housekeeping gene (GAPDH) and normalised to their respective iPSCs (n = 1 for each line, 3 technical replicates). Error bars indicate the Standard Error of Mean. **(E)** qRT-PCR for TH, SNCA, NURR1, DAPT and DAT at day 65. mRNA values are relative to the housekeeping gene and normalised to the corresponding iPSCs (n = 3-5 per line). **(F)** Cropped immunoblot of total WDR45 and beta actin protein expression at Day 11, and relevant quantification. n=3-4 biological replicates for each line. Error bars represent the Standard Error of Mean. Statistics were calculated using ANOVA. Abbreviations: EBs= embryoid bodies. FC= fold change

**Supplementary Figure 3.**
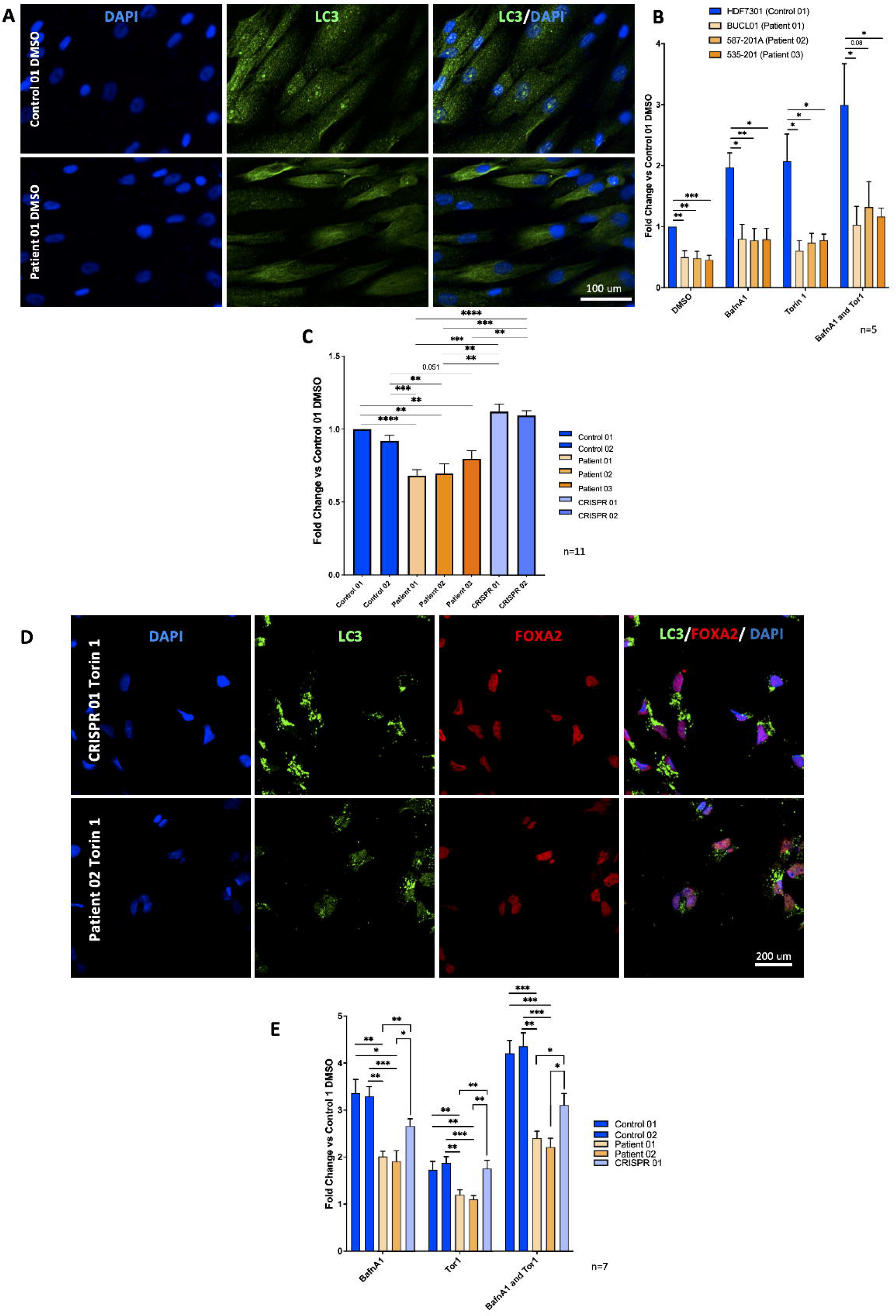
RNASeq at Day 65 of differentiation. **(A)** List of differentially expressed genes when comparing Patient 01, Patient 02, Patient 03 versus Control 01, Control 02, CRISPR 01 and CRISPR 02 mDA neurons. **(B)** ClueGO analysis of GO terms enrichment of differentially expressed genes, showing pie charts for cellular component (CC), and molecular function (MF). **(C)** Volcano plots of differentially expressed genes when comparing Patient 02 and corresponding CRISPR line (CRISPR 01), as well as Patient 03 versus corresponding CRISPR line (CRISPR 02). The top 40 genes (as per lowest p-values) are labelled. Right: GO Term and KEGG pathway enrichment analysis depicting intracellular pathways jointly corrected in both Patients 02 and 03, when compared to CRISPR 01 and 02. **(D)** List of intracellular pathways and genes corrected in both Patients 02 and 03, when compared to CRISPR 01 and 02. **(E)** List of differentially expressed genes and involved pathways when comparing Patient 02, versus CRISPR 01 (Patient 02 Corrected) and Patient 02 Torin 1- and Digoxin-treated mDA neurons. **(F)** ClueGO analysis of GO terms enrichment of differentially expressed genes, showing pie charts for cellular component (CC), and molecular function (MF). n=3 for all lines, median TPM values analysed. Network graph nodes represent GO terms (the most significant are named) and edges indicate shared genes between GO terms. Functional groups of GO terms are indicated by the same colour. Pie charts show the percentages of each functional group representation. GO functional groups exhibiting statistically significant differences (p< 0.05) are shown.

**Supplementary Figure 4.**
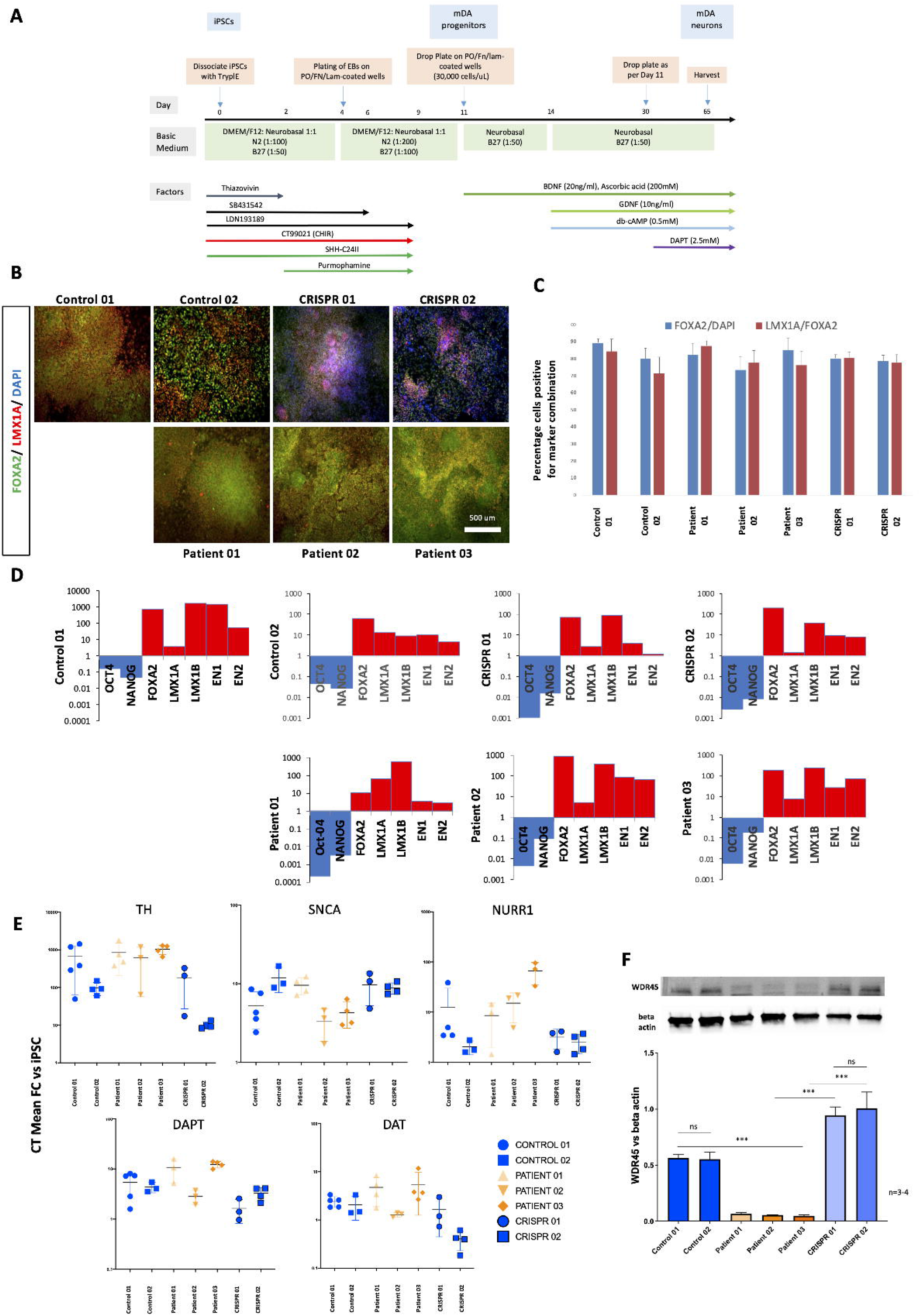
Defective autophagy flux in BPAN cells. (A) Patient and control fibroblasts imaged after 3-hour treatments with DMSO, autophagy flux inducers (Torin 1) and/ or inhibitors (Bafilomycin A1). Representative images. Cells were plated in 96-well plates at a density of 15,000 cells/well. n=5 biological replicates for each line. For each biological replicate, all lines were seeded on the same 96-well plate. (B) Quantification of LC3 puncta/ nuclei in control and patient-derived fibroblasts. For statistical analysis, the Student’s unpaired two tailed t-test was used. Error bars represent the Standard Error of Mean. (C) Quantification of LC3 puncta/ nuclei in control and patient-derived neuronal progenitors, at basal (DMSO-treated) conditions. Experiment identical to the one depicted in Fig. 3A-B, but with more independent biological replicates (n=11). Additional replicates enhance the statistical significance of previous findings. For statistical analysis, the Student’s unpaired two tailed t-test was used. Error bars represent the Standard Error of Mean. (D) Day 11 ventral progenitors imaged after 3-hour autophagy flux induction or inhibition. Representative images. Cells were plated in 96-well plates at a density of 15,000 cells/well. n=7 independent differentiations/ biological replicates for each line. For each biological replicate, all 5 lines had the same start date of differentiation and were seeded on the same 96-well plate. (E) Quantification of LC3 puncta/ nuclei in control and patient-derived neurons. For statistical analysis, the Student’s unpaired two tailed t-test was used. Error bars represent the Standard Error of Mean.

## Supplementary Tables

**Supplementary Table 1.** Fibroblast and corresponding iPSC lines used. After characterisation of pluripotency, one iPSC clone from each line was used for downstream differentiations. The first patient line (Patient 01, BPAN07) carries a splice site mutation that leads to aberrant splicing and an early stop codon. Alignment of wild type & WDR45 c.344+2T>A amino acid sequences shows premature truncation of the protein by 246 amino acids with the inclusion of 2 aberrant residues (arginine and Alanine); p.(Ile116Argfs*3) (data not shown). The other two patient lines (587-02 and 535-02) harbour nonsense pathogenic mutations leading to an early stop codon. In the isogenic controls R7-72 and R234-68, disease-causing mutations (in Patients 02 and 03, respectively) were corrected using CRISPR/Cas9-mediated genome editing (Supplementary Figure 1). Age-matched healthy control fibroblasts HDF-7301 were collected from the MRC Centre for Neuromuscular Disorders Biobank. Patient fibroblast line BUCL01 was ascertained from the University College London (UCL) Great Ormond Street Institute of Child Health (UCL GOS ICH), London, UK. Control fibroblast line 582-202 and patient lines 587-201A and 535-201 were obtained from Oregon Health and Science University (OHSU), Portland, Oregon, USA. Patient BUCL01 and control HDF-7301 fibroblasts were reprogrammed into iPSC at UCL GOS ICH, while control 582-202 and patients 587-201A and 535-201 fibroblasts at the Wellcome Trust-Medical Research Council Cambridge Stem Cell Institute (Anne McLaren Laboratory for Regenerative Medicine, Cambridge, UK). Lines 587-02 and 535-02 (as well as the isogenic controls R7-72 and R234-68) were initially plated on Vitronectin XF (Stemcell Technologies)-coated plates and cultured in TeSR-E8 (StemCell Technologies). These lines were subsequently transferred to Matrigel/ mTeSR1 culture conditions.

| Fibroblast Identifier | iPSC clone used after characterisation | WDR45 mutation | Protein Effect | Age at biopsy | Gender |
| --- | --- | --- | --- | --- | --- |
| HDF7301 | HDF7301-05 (Control 01) | Healthy Control | - | 13y | F |
| 582-202 | 582-06 (Control 02) | Healthy Control | - | 1y10mo | M |
| BUCL01 | BPAN07 (Patient 01) | c.344+2T>A | p.(Ile116Argfs*3) | 5y | F |
| 587-201A | 587-02 (Patient 02) | c.19C>T | p.Arg7* | 19y8mo | M |
| 535-201 | 535-02 (Patient 03) | c.700C>T | p.Arg234* | 6y4mo | F |
|  | R7-72 (CRISPR 01, Patient 02 Corrected) | Substitution of c.19C>T with 'wild-type' sequence | Normal length | 19y8mo | M |
|  | R234-68 (CRISPR 02, Patient 03 Corrected) | Substitution of c.19C>T with 'wild-type' sequence | Normal length | 6y4mo | F |

**Supplementary Table 2.**
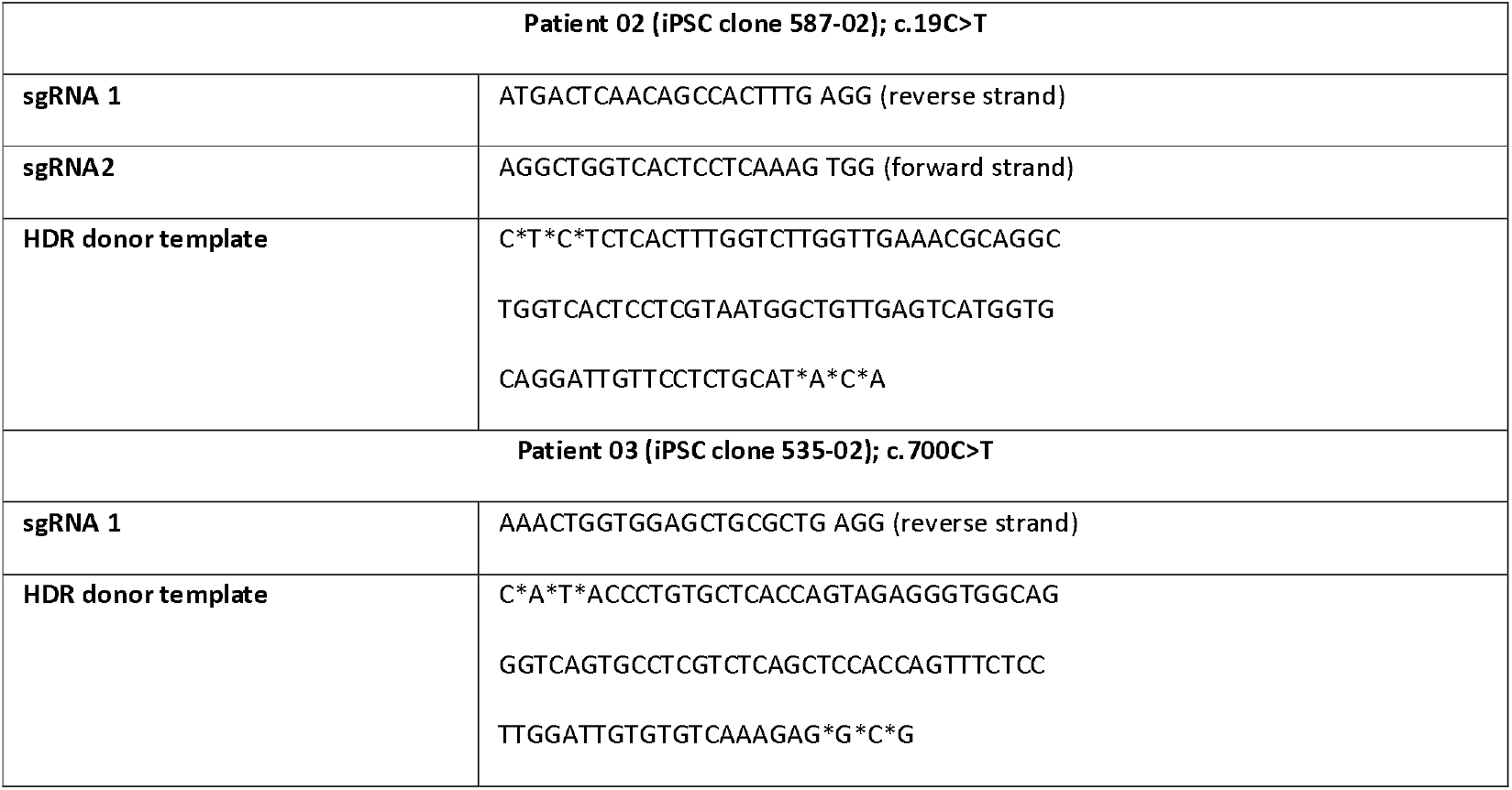
sgRNA and HDR donor templates used for CRISPR/Cas9 genome editing in patient lines.

**Supplementary Table 3.**
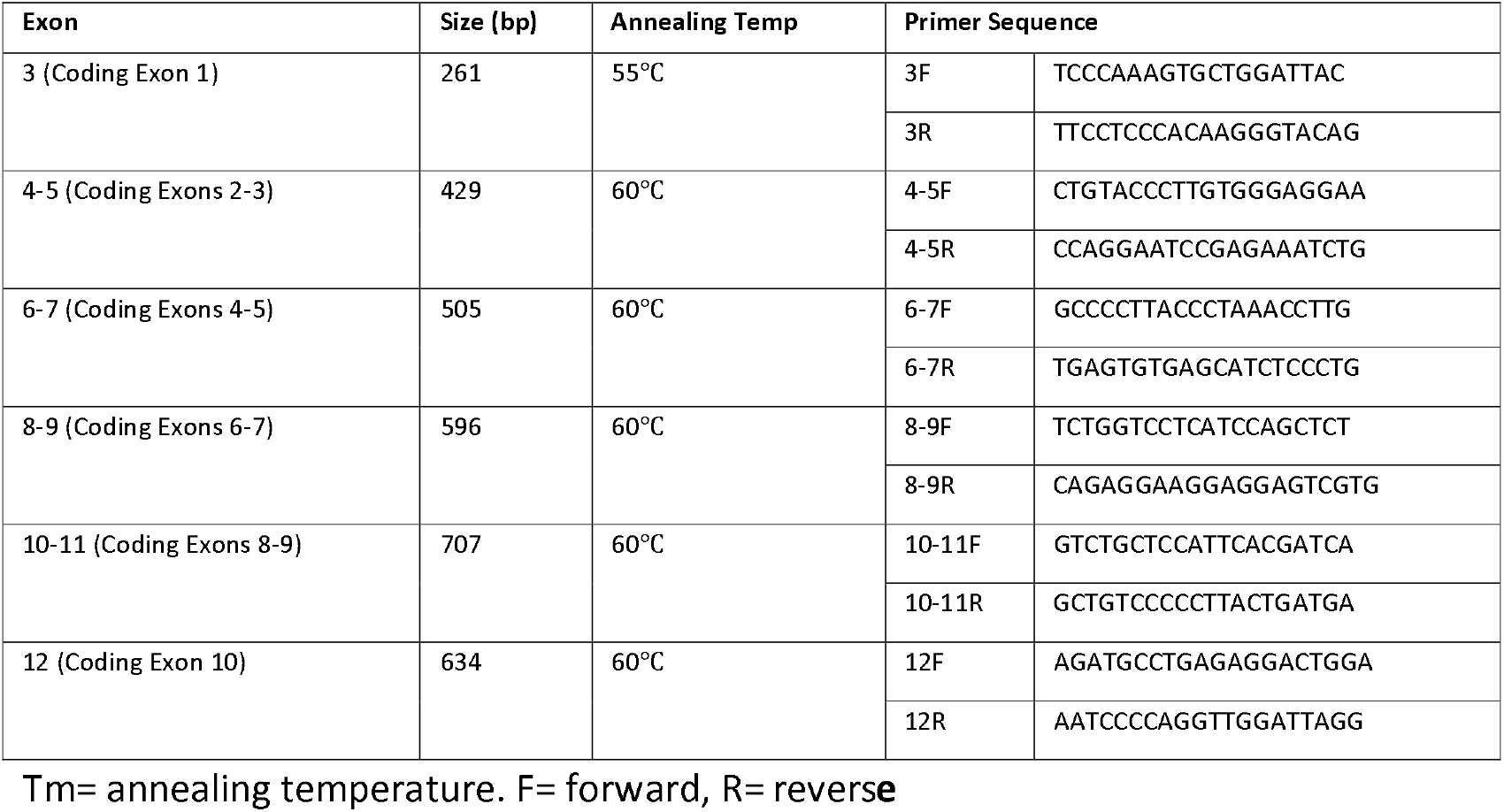
PCR primers for WDR45 gene sequencing.

**Supplementary Table 4.**
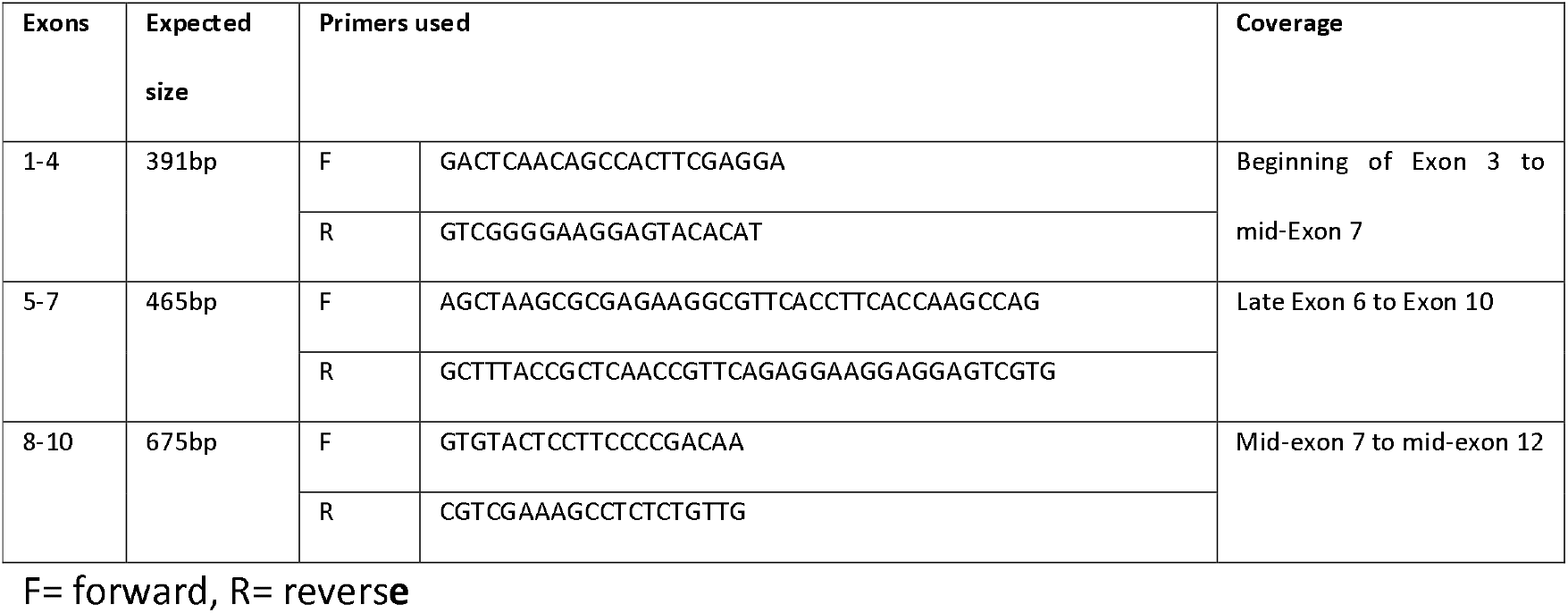
Primers used for WDR45 cDNA sequencing.

**Supplementary Table 5.**
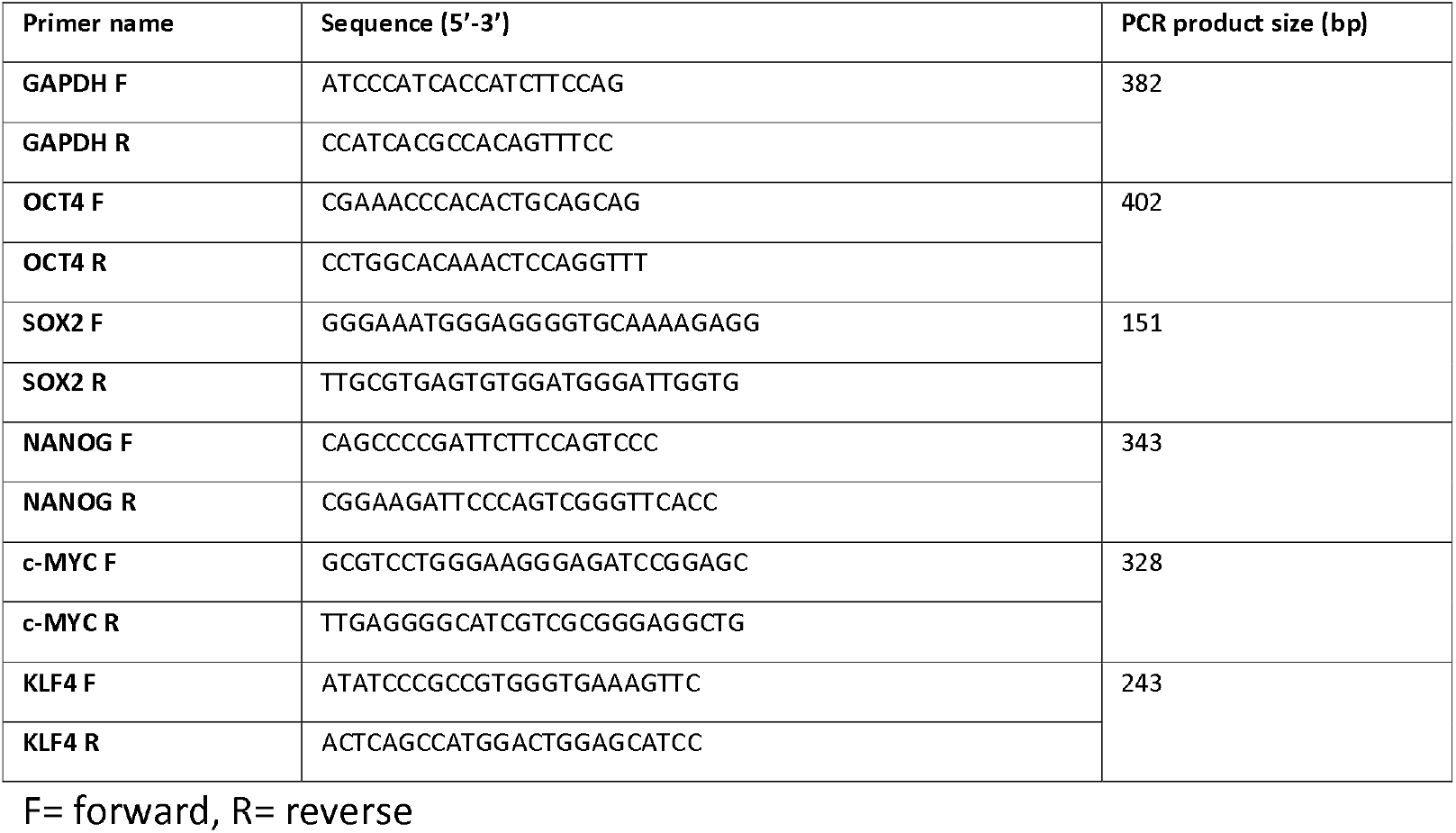
Primer pairs for detection of pluripotency marker expression via RT PCR.

**Supplementary Table 6.**
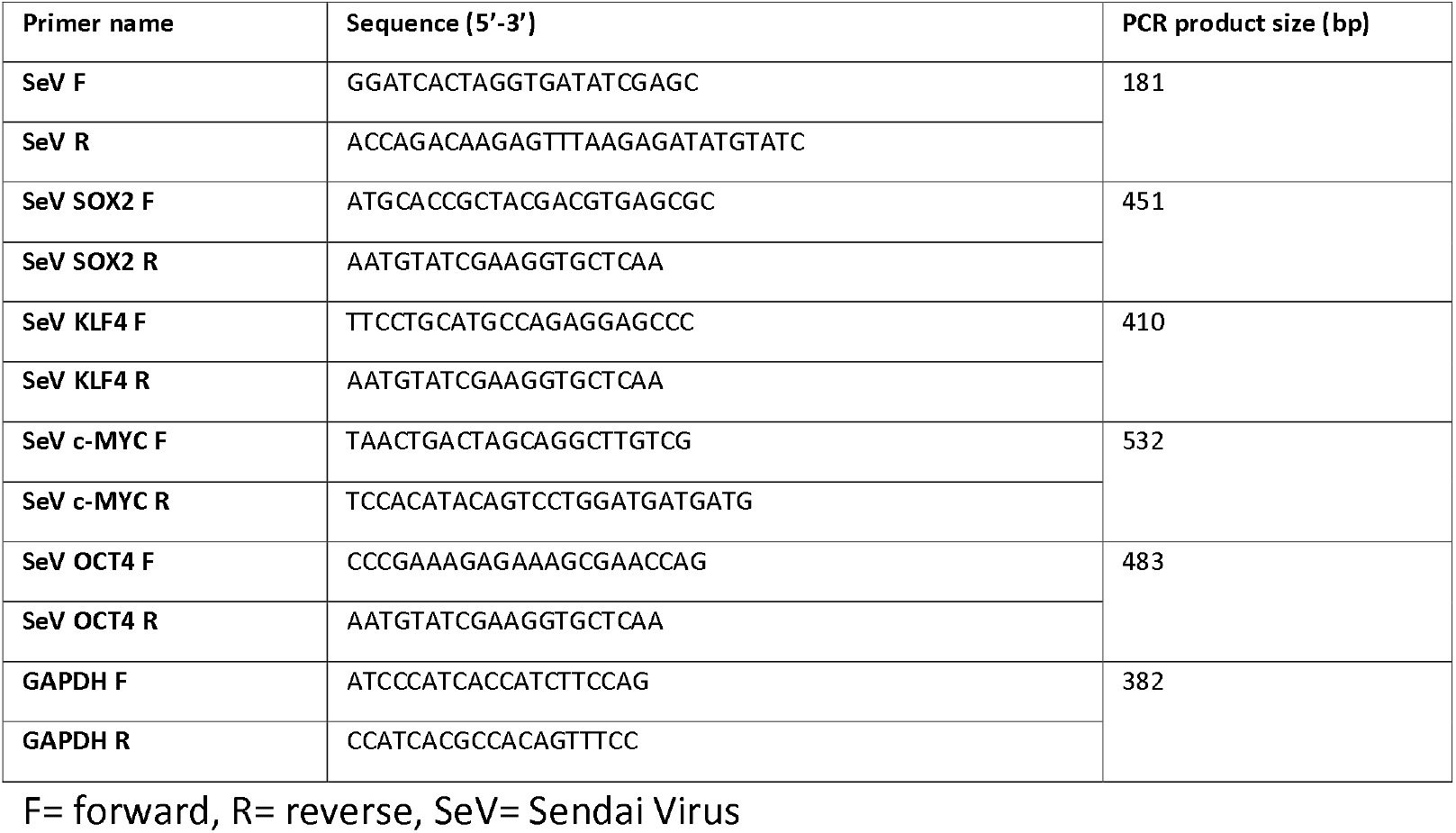
Primers used for Sendai Virus Clearance-related RT PCR experiments.

**Supplementary Table 7.**
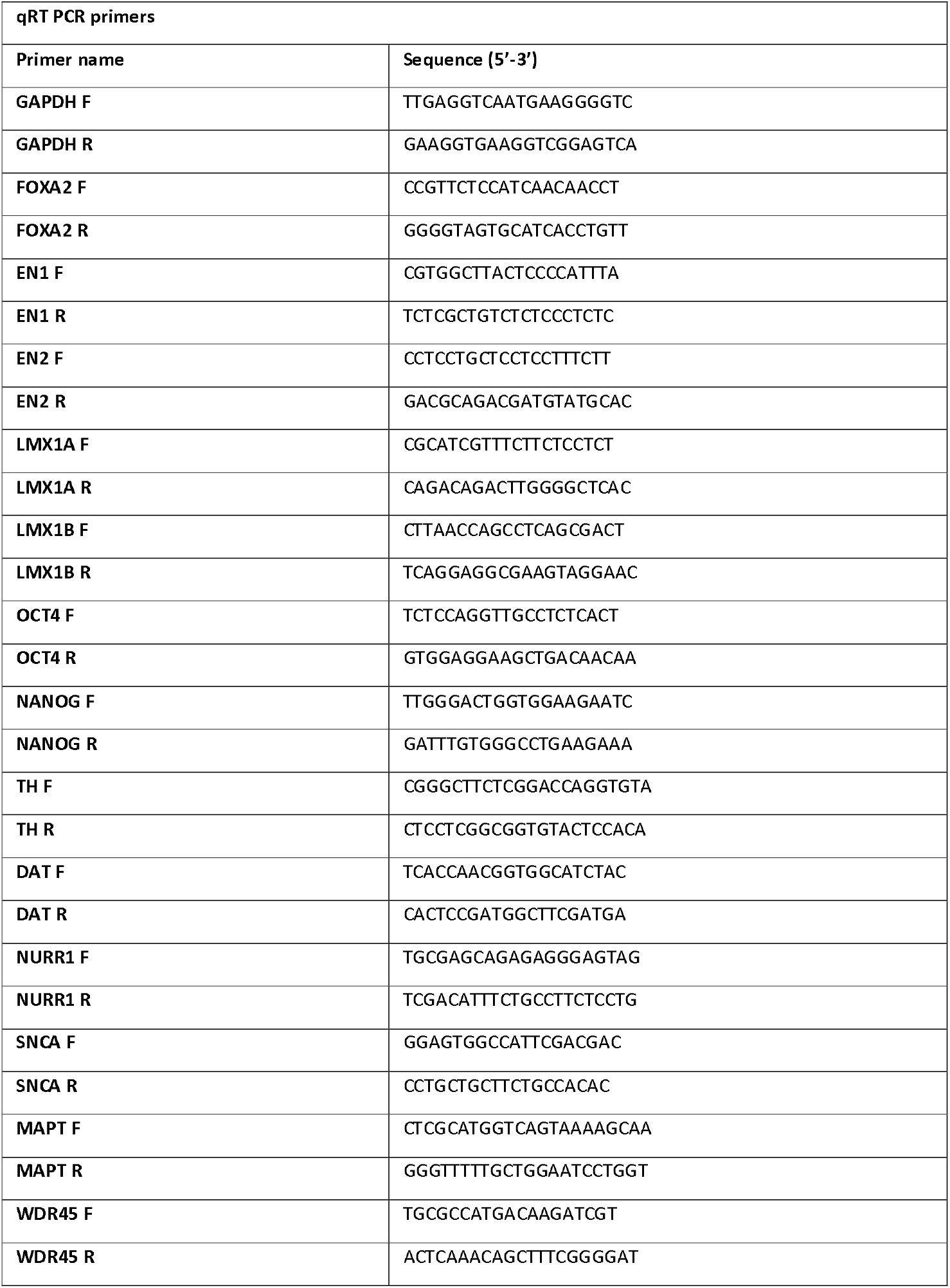
qRT PCR primers used for Day 11 and Day 65 characterisation.

**Supplementary Table 8.** Primary and corresponding secondary antibodies used for immunofluorescence and western blotting experiments.

| Western Blot Antibodies |  |  |
| --- | --- | --- |
| Primary Antibody | Company/ Catalogue Number | Dilution |
| Beta Actin mouse monoclonal antibody (clone AC-15) | Sigma (A1978) | 1:4,000 |
| LC3B rabbit polyclonal antibody | Sigma (L7543) | 1:1,000 |
| WDR45 rabbit monoclonal antibody | Gift from Professor Sharon Tooze's laboratory, Francis Crick Institute, London, UK | 1:250 |
| WDR45 rabbit polyclonal antibody | Proteintech (19194-1-AP) | 1:2,000 |
| Secondary Antibody | Company/ Catalogue Number | Dilution |
| Anti-mouse IgG, HRP-linked Antibody | Cell signalling technology (7076) | 1:5,000 |
| Anti-rabbit IgG, HRP-linked Antibody | Cell signalling technology (7074) | 1:5,000 |
| Immunofluorescence Antibodies |  |  |
| Primary Antibody | Company/ Catalogue Number | Dilution |
| FOXA2 mouse monoclonal antibody | BD Pharmigen (561580) | 1:500 |
| LMX1A rabbit polyclonal antibody | Millipore (AB10533) | 1:2,000 |
| MAP2 Mouse monoclonal antibody | Sigma | 1:400 |
| NANOG Mouse monoclonal antibody | Millipore | 1:500 |
| OCT4 Mouse monoclonal antibody | Santa Cruz | 1:50 |
| SMA Rabbit monoclonal antibody | Abcam | 1:100 |
| SOX17 Goat polyclonal antibody | R&D Systems | 1:200 |
| TH Chicken polyclonal antibody | Aves | 1:400 |
| TRA-1-60 Mouse monoclonal antibody | Santa Cruz | 1:200 |
| TRA-1-81 Mouse monoclonal antibody | Millipore | 1:200 |
| TUJ1 Mouse monoclonal antibody | BioLegend | 1:400 |
| Secondary Antibody | Company/ Catalogue Number | Dilution |
| Alexa Fluor 488 Donkey Anti-Goat IgG | ThermoFisher (A-11055) | 1:400 |
| Alexa Fluor 488 Goat Anti-Mouse IgG | ThermoFisher (A-11001) | 1:400 |
| Alexa Fluor 488 Goat Anti-Rabbit IgG | ThermoFisher (A-11008) | 1:400 |
| Alexa Fluor 594 Goat Anti-Chicken IgG | ThermoFisher (A-11042) | 1:400 |
| Alexa Fluor 594 Goat Anti-Mouse IgM | Thermofisher (A-21044) | 1:400 |
| Alexa Fluor 594 Goat Anti-Rabbit IgG | ThermoFisher (A-11012) | 1:400 |
| High Content Immunofluorescence Antibodies |  |  |
| Primary Antibody | Company/ Catalogue Number | Dilution |
| LC3B rabbit monoclonal antibody (clone D11) | Cell Signalling Technology (3868S) | 1:200 |
| P62/SQSTM1 rabbit polyclonal antibody | Sigma (P0067) | 1:1,000 |
| Secondary Antibody | Company/ Catalogue Number | Dilution |
| Alexa Fluor 488 Goat Anti-Mouse IgG | ThermoFisher (A-11001) | 1:400 |
| Alexa Fluor 488 Goat Anti-Rabbit IgG | ThermoFisher (A-11008) | 1:400 |

**Supplementary Table 9.** Hits from Prestwick screen with the 200 highest z-scores.

| Hit | Source plate | Well | Z score | Compound |
| --- | --- | --- | --- | --- |
| 1 | 14 | A10 | 253.64 | Oxyphenbutazone |
| 2 | 6 | D09 | 152.67 | Doxorubicin hydrochloride |
| 3 | 14 | C09 | 107.82 | Thiethylperazine dimaleate |
| 4 | 10 | F08 | 93.28 | Alexidine dihydrochloride |
| 5 | 7 | A08 | 82.75 | Daunorubicin hydrochloride |
| 6 | 9 | D07 | 60.76 | Beta-Escin |
| 7 | 15 | F02 | 57 | Topotecan |
| 8 | 9 | B07 | 53.24 | Lanatoside C |
| 9 | 12 | A04 | 46.93 | Digoxigenin |
| 10 | 12 | E06 | 39.85 | Thonzonium bromide |
| 11 | 11 | B02 | 32.34 | Auranofin |
| 12 | 8 | H11 | 25.03 | Sertindole |
| 13 | 16 | G02 | 24.93 | Epirubicin hydrochloride |
| 14 | 16 | B05 | 24.49 | Tegaserod maleate |
| 15 | 9 | G09 | 24.39 | Benzethonium chloride |
| 16 | 6 | D08 | 21.69 | Digoxin |
| 17 | 5 | G06 | 21.09 | Mitoxantrone dihydrochloride |
| 18 | 9 | G06 | 19.4 | Methyl benzethonium chloride |
| 19 | 7 | G11 | 18.22 | Parthenolide |
| 20 | 6 | D07 | 17.8 | Digitoxigenin |
| 21 | 5 | H08 | 15.89 | Clomiphene citrate (Z,E) |
| 22 | 9 | A11 | 14.96 | Pinaverium bromide |
| 23 | 4 | E07 | 14.67 | Perhexiline maleate |
| 24 | 13 | C07 | 14.2 | Proscillaridin A |
| 25 | 1 | H09 | 13.55 | Thioridazine hydrochloride |
| 26 | 6 | A10 | 13.41 | Amiodarone hydrochloride |
| 27 | 2 | F07 | 12.87 | Astemizole |
| 28 | 7 | E03 | 11.92 | Thiostrepton |
| 29 | 13 | H11 | 11.5 | Pyrvinium pamoate |
| 30 | 4 | H09 | 10.16 | Quinacrine dihydrochloride hydrate |
| 31 | 5 | F06 | 10.13 | Methiothepin maleate |
| 32 | 10 | G08 | 9.97 | Merbromin |
| 33 | 9 | A07 | 9.16 | Oxiconazole Nitrate |
| 34 | 2 | E07 | 9.1 | Mefloquine hydrochloride |
| 35 | 2 | G07 | 8.59 | Tamoxifen citrate |
| 36 | 5 | E09 | 8.14 | Bepiridil hydrochloride |
| 37 | 14 | A06 | 7.55 | Sertaconazole nitrate |
| 38 | 7 | H11 | 7.49 | Prenylamine lactate |
| 39 | 13 | A11 | 7.18 | Indatraline hydrochloride |
| 40 | 5 | G07 | 6.77 | GBR 12909 dihydrochloride |
| 41 | 16 | E05 | 6.53 | Benzoxiquine |
| 42 | 2 | G04 | 6.11 | Chlorhexidine |
| 43 | 4 | H04 | 6.06 | Trifluoperazine dihydrochloride |
| 44 | 6 | F08 | 5.67 | Meclozine dihydrochloride |
| 45 | 3 | D11 | 5.59 | Camptothecin (S, +) |
| 46 | 4 | C11 | 5.49 | Fendiline hydrochloride |
| 47 | 11 | A11 | 5.47 | Sulconazole nitrate |
| 48 | 10 | C08 | 5.44 | Aprepitant |
| 49 | 14 | C11 | 5.42 | Vorinostat |
| 50 | 9 | B11 | 5.37 | Avermectin B1a |
| 51 | 5 | E05 | 4.47 | Metergoline |
| 52 | 13 | H02 | 4.27 | Halofantrine hydrochloride |
| 53 | 9 | C07 | 4.17 | Nisoldipine |
| 54 | 4 | H11 | 4.16 | Fluphenazine dihydrochloride |
| 55 | 1 | H12 | 4 | Positive Control |
| 56 | 15 | H11 | 3.94 | Hexachlorophene |
| 57 | 5 | C09 | 3.91 | Chlorprothixene hydrochloride |
| 58 | 10 | G12 | 3.51 | Positive Control |
| 59 | 7 | G02 | 3.43 | Ciclopirox ethanolamine |
| 60 | 4 | G05 | 3.43 | Econazole nitrate |
| 61 | 12 | A12 | 3.27 | Positive Control |
| 62 | 1 | H07 | 3.13 | Dibucaine |
| 63 | 2 | H12 | 3.12 | Positive Control |
| 64 | 4 | H03 | 3.02 | Flunarizine dihydrochloride |
| 65 | 11 | G12 | 2.91 | Positive Control |
| 66 | 1 | A12 | 2.85 | Positive Control |
| 67 | 11 | G07 | 2.84 | Azacytidine-5 |
| 68 | 3 | A02 | 2.84 | Isoniazid |
| 69 | 2 | E05 | 2.83 | Tioconazole |
| 70 | 13 | F05 | 2.79 | Sertraline |
| 71 | 2 | G12 | 2.7 | Positive Control |
| 72 | 1 | F12 | 2.69 | Positive Control |
| 73 | 11 | F12 | 2.68 | Positive Control |
| 74 | 1 | A02 | 2.66 | Azaguanine-8 |
| 75 | 1 | D12 | 2.6 | Positive Control |
| 76 | 10 | F12 | 2.59 | Positive Control |
| 77 | 10 | H12 | 2.56 | Positive Control |
| 78 | 6 | B05 | 2.56 | Amlodipine |
| 79 | 16 | G05 | 2.55 | Lomerizine hydrochloride |
| 80 | 11 | G03 | 2.55 | Raloxifene hydrochloride |
| 81 | 5 | H10 | 2.52 | Prochlorperazine dimaleate |
| 82 | 5 | G04 | 2.5 | Nicardipine hydrochloride |
| 83 | 5 | H07 | 2.5 | Etoposide |
| 84 | 2 | D12 | 2.49 | Positive Control |
| 85 | 14 | H12 | 2.48 | Positive Control |
| 86 | 13 | F12 | 2.47 | Positive Control |
| 87 | 6 | A11 | 2.46 | Amphotericin B |
| 88 | 1 | C12 | 2.45 | Positive Control |
| 89 | 16 | H11 | 2.44 | Cefprozil |
| 90 | 11 | A12 | 2.43 | Positive Control |
| 91 | 16 | H10 | 2.3 | Desonide |
| 92 | 14 | H06 | 2.29 | Ronidazole |
| 93 | 1 | G12 | 2.27 | Positive Control |
| 94 | 4 | H10 | 2.26 | Clofilium tosylate |
| 95 | 1 | F08 | 2.25 | Benoxinate hydrochloride |
| 96 | 11 | G06 | 2.24 | Simvastatin |
| 97 | 5 | F12 | 2.2 | Positive Control |
| 98 | 10 | B11 | 2.19 | Ebselen |
| 99 | 10 | E12 | 2.16 | Positive Control |
| 100 | 3 | B02 | 2.14 | Tranexamic acid |
| 101 | 12 | A03 | 2.13 | Bemegride |
| 102 | 16 | H12 | 2.13 | Positive Control |
| 103 | 11 | H12 | 2.11 | Positive Control |
| 104 | 16 | G12 | 2.1 | Positive Control |
| 105 | 12 | A08 | 2.09 | Oxybenzone |
| 106 | 7 | H05 | 2.08 | Tolazamide |
| 107 | 1 | B12 | 2.07 | Positive Control |
| 108 | 4 | F12 | 2.06 | Positive Control |
| 109 | 7 | D01 | 2.04 | Positive Control |
| 110 | 10 | A10 | 2.02 | Sulfanilamide |
| 111 | 12 | A09 | 2 | Promethazine hydrochloride |
| 112 | 5 | C12 | 1.99 | Positive Control |
| 113 | 12 | B03 | 1.98 | Flubendazol |
| 114 | 2 | C12 | 1.97 | Positive Control |
| 115 | 10 | B12 | 1.87 | Positive Control |
| 116 | 8 | H07 | 1.86 | Penciclovir |
| 117 | 11 | B12 | 1.86 | Positive Control |
| 118 | 7 | H04 | 1.86 | Pentamidine isethionate |
| 119 | 12 | A01 | 1.83 | Positive Control |
| 120 | 6 | B10 | 1.82 | Bisacodyl |
| 121 | 1 | F04 | 1.8 | Triflupromazine hydrochloride |
| 122 | 5 | C08 | 1.79 | Thioguanosine |
| 123 | 15 | H12 | 1.77 | Positive Control |
| 124 | 9 | A03 | 1.76 | Gefitinib |
| 125 | 16 | H08 | 1.75 | Triclabendazole |
| 126 | 8 | H12 | 1.74 | Positive Control |
| 127 | 7 | A05 | 1.73 | Benperidol |
| 128 | 13 | D12 | 1.71 | Positive Control |
| 129 | 2 | E12 | 1.71 | Positive Control |
| 130 | 4 | H12 | 1.71 | Positive Control |
| 131 | 14 | E03 | 1.69 | (-)-Eseroline fumarate salt |
| 132 | 13 | D09 | 1.69 | Zuclopenthixol dihydrochloride |
| 133 | 12 | A07 | 1.69 | Cloquinol |
| 134 | 16 | F11 | 1.67 | Ritonavir |
| 135 | 16 | E01 | 1.67 | Positive Control |
| 136 | 14 | F12 | 1.66 | Positive Control |
| 137 | 1 | H08 | 1.66 | Prednisone |
| 138 | 15 | H09 | 1.65 | Sildenafil |
| 139 | 16 | H04 | 1.64 | Raltitrexed |
| 140 | 16 | H03 | 1.62 | Pemetrexed disodium |
| 141 | 7 | H03 | 1.6 | Selegiline hydrochloride |
| 142 | 3 | B01 | 1.59 | Positive Control |
| 143 | 12 | D12 | 1.59 | Positive Control |
| 144 | 11 | A10 | 1.59 | Dichlorphenamide |
| 145 | 16 | D02 | 1.58 | Aminacrine |
| 146 | 9 | H07 | 1.57 | Pramoxine hydrochloride |
| 147 | 2 | B11 | 1.57 | Nocodazole |
| 148 | 11 | D12 | 1.57 | Positive Control |
| 149 | 7 | C12 | 1.57 | Positive Control |
| 150 | 4 | G12 | 1.55 | Positive Control |
| 151 | 6 | H07 | 1.54 | Primaquine diphosphate |
| 152 | 12 | A10 | 1.54 | Diacerein |
| 153 | 1 | C07 | 1.53 | Cimetidine |
| 154 | 6 | H06 | 1.53 | Brompheniramine maleate |
| 155 | 10 | A09 | 1.51 | Sulfadimethoxine |
| 156 | 1 | G10 | 1.51 | Acebutolol hydrochloride |
| 157 | 1 | F11 | 1.5 | Tolazoline hydrochloride |
| 158 | 11 | H06 | 1.5 | Reserpine |
| 159 | 10 | H09 | 1.49 | Dienestrol |
| 160 | 13 | E12 | 1.48 | Positive Control |
| 161 | 16 | H07 | 1.48 | Milnacipran hydrochloride |
| 162 | 11 | E12 | 1.47 | Positive Control |
| 163 | 10 | H10 | 1.47 | Pridinol methanesulfonate salt |
| 164 | 16 | H05 | 1.47 | Ceftibuten |
| 165 | 6 | H09 | 1.46 | Felodipine |
| 166 | 10 | A04 | 1.46 | Phenethicillin potassium salt |
| 167 | 10 | H11 | 1.45 | Amrinone |
| 168 | 7 | D12 | 1.45 | Positive Control |
| 169 | 8 | H05 | 1.43 | Tomoxetine hydrochloride |
| 170 | 10 | G10 | 1.43 | Drofenine hydrochloride |
| 171 | 13 | H06 | 1.42 | Molindone hydrochloride |
| 172 | 10 | G03 | 1.42 | Podophyllotoxin |
| 173 | 16 | D12 | 1.42 | Positive Control |
| 174 | 10 | D12 | 1.42 | Positive Control |
| 175 | 2 | H11 | 1.41 | Gentamicine sulfate |
| 176 | 12 | C12 | 1.41 | Positive Control |
| 177 | 15 | H08 | 1.4 | Rivastigmine |
| 178 | 8 | H10 | 1.4 | Etoricoxib |
| 179 | 11 | A09 | 1.39 | Furazolidone |
| 180 | 1 | G08 | 1.39 | Miconazole |
| 181 | 5 | D12 | 1.39 | Positive Control |
| 182 | 15 | G12 | 1.38 | Positive Control |
| 183 | 1 | H11 | 1.37 | Trimethobenzamide hydrochloride |
| 184 | 8 | H09 | 1.37 | Dexfenfluramine hydrochloride |
| 185 | 6 | A05 | 1.36 | Idobenone |
| 186 | 1 | H06 | 1.35 | Adiphenine hydrochloride |
| 187 | 6 | B01 | 1.34 | Positive Control |
| 188 | 16 | E12 | 1.33 | Positive Control |
| 189 | 13 | C12 | 1.33 | Positive Control |
| 190 | 1 | E12 | 1.33 | Positive Control |
| 191 | 5 | F07 | 1.33 | Clofazimine |
| 192 | 10 | C09 | 1.32 | Monensin sodium salt |
| 193 | 13 | H07 | 1.32 | Alcuronium chloride |
| 194 | 13 | G04 | 1.32 | Isometheptene mucate |
| 195 | 8 | H04 | 1.3 | Thiorphan |
| 196 | 13 | C10 | 1.3 | Chlormadinone acetate |
| 197 | 3 | A03 | 1.3 | Pentylentetrazole |
| 198 | 1 | F09 | 1.28 | Oxethazaine |
| 199 | 14 | H09 | 1.26 | Cefepime hydrochloride |
| 200 | 11 | A07 | 1.25 | Trimipramine maleate salt |

**Supplementary Table 10.** ATG differential gene expression in Day 65 mDA Patient 02 cultures after CRISPR correction and compound treatments. Genes marked in blue show consistent under- or over-expression when comparing the Patient 02 untreated mDA line profiles with the corresponding CRISPR-corrected and compound-treated ones. Many known ATGs were interrogated. P-value and fold change cut-offs were not applied for this analysis; however, some genes have significant p- and fold change values in different conditions. ATG= autophagy-related gene, FC= fold change.

| ATG | Patient 02 vs (Controls 01 and 02) |  | CRISPR 01 vs Patient 02 |  | Digoxin-treated line vs Patient 02 |  | Torin 1-treated vs Patient 02 |  |
| --- | --- | --- | --- | --- | --- | --- | --- | --- |
|  | FC | P-Value | FC | P-Value | FC | P-Value | FC | P-Value |
| AMBRA1 | 0.43706262 (down) | 6.14E-04 | 0.7145338 (down) | 0.16574542 | 1.2240671 (up) | 0.487703 | 0.8595378 (down) | 0.7000872 |
| ATG12 | 1.0688772 (no change) | 0.792146 | 1.1249504 (up) | 0.10514027 | 0.92068404 (down) | 0.61909074 | 1.168334 (up) | 0.33339965 |
| ATG13 | 0.9291793 (down) | 0.7693057 | 0.93510157 (down) | 0.5057616 | 0.4419558 (down) | 0.0680989 | 0.90944344 (down) | 0.42617658 |
| ATG14 | 1.1413882 (up) | 0.5587088 | Not mapped | Not mapped | 0.6088769 (down) | 0.17958279 | Not mapped | Not mapped |
| ATG16L1 | 0.59600216 (down) | 0.31001356 | 1.0367168 (no change) | 0.9407053 | 2.9104712 (up) | 0.00144182 | 1.3612424 (up) | 0.54965794 |
| ATG16L2 | 0.71721536 (down) | 0.010059934 | Not mapped | Not mapped | Not mapped | Not mapped | 1.1957258 (up) | 0.6599298 |
| ATG2A | 0.9201323 (down) | 0.8116482 | 1.986146 (up) | 0.16611445 | 1.2601732 (up) | 0.6257956 | 1.1654925 (up) | 0.7045777 |
| ATG2B | 0.81731015 (down) | 0.24680214 | 1.335941 (up) | 0.35303855 | 2.0343752 (up) | 0.04702732 | 1.1501077 (up) | 0.5015331 |
| ATG3 | 0.84620136 (down) | 0.44752243 | Not mapped | Not mapped | 1.0925838 (no change) | 0.8455659 | 0.81203955 (down) | 0.6288875 |
| ATG4B | 1.6198123 (up) | 0.5479161 | 0.7977216 (down) | 0.29556963 | 0.5249194 (down) | 0.4338038 | 0.3615233 (down) | 0.00814244 |
| ATG5 | 2.085391 (up) | 0.01342523 | Not mapped | Not mapped | 0.5981643 (down) | 0.27420387 | 0.73430216 (down) | 0.37415075 |
| BECN1 | 4.088446 (up) | 0.19371912 | 0.43691146 (down) | 0.5124709 | 0.09202237 (down) | 0.21182562 | 0.15318666 (down) | 0.28554642 |
| EPG5 | 0.891338 (down) | 0.8618909 | 1.0892509 (no change) | 0.6674719 | 2.3882854 (up) | 0.17077148 | 1.0209918 (no change) | 0.8568484 |
| GABARAP | 1.0952392 (no change) | 0.78651476 | 0.89669627 (down) | 0.6855389 | 0.60620856 (down) | 0.08376524 | 1.1691217 (up) | 0.5665487 |
| GABARAPL1 | 0.8820572 (down) | 0.5491498 | 1.0306478 (no change) | 0.85970575 | 1.750449 (up) | 0.12943085 | 1.3634212 (up) | 0.24656725 |
| GABARAPL2 | 0.9287938 (down) | 0.82963806 | 0.8410031 (down) | 0.62539303 | 0.75546306 (down) | 0.20855312 | 1.3366152 (up) | 0.37566233 |
| MAP1LC3A | 0.8614579 (down) | 0.6068979 | 1.5705838 (up) | 0.20977472 | 1.4284861 (up) | 0.01311642 | 1.3044173 (up) | 0.02771006 |
| MAP1LC3B | 0.9986874 (down) | 0.99608976 | 0.7477211 (down) | 0.19170646 | 1.5358086 (up) | 0.48995838 | 0.87073094 (down) | 0.72981685 |
| RAB24 | 1.0563483 (no change) | 0.7662744 | Not mapped | Not mapped | Not mapped | Not mapped | Not mapped | Not mapped |
| RAB7A | 0.91218925 (down) | 0.57352966 | 0.97811717 (down) | 0.91971004 | 1.3447509 (up) | 0.07130751 | 1.143196 (up) | 0.4439764 |
| ULK1 | 0.6885398 (down) | 0.32386762 | 2.6435022 (up) | 0.00500735 | 0.46201056 (down) | 0.03606971 | 2.4438975 (up) | 0.00704683 |
| ULK2 | 1.6028628 (up) | 0.43187773 | 1.4053199 (up) | 0.04052436 | 0.5006925 (down) | 0.01474166 | 1.067213 (no change) | 0.8066371 |
| UVRAG | 0.16716082 (down) | 1.77E-05 | 2.3673134 (up) | 0.05409699 | 3.923307 (up) | 0.00618942 | 2.1328526 (up) | 0.03366591 |
| WIPI1 | 1.1785417 (up) | 0.62346715 | 0.43872187 (down) | 0.01289181 | 0.5140383 (down) | 0.01987111 | 1.1200486 (up) | 0.5993699 |
| WIPI2 | 0.71252674 (down) | 0.025780724 | 0.6941722 (down) | 0.19979995 | Not mapped | Not mapped | 1.2907548 (up) | 0.11014623 |
| WIPI3 | 0.8958507 (down) | 0.6354098 | 1.0138819 (no change) | 0.96299773 | 0.80224526 (down) | 0.4812425 | 0.9755141 (down) | 0.93992513 |
| WDR45 | 2.1794524 (up) | 0.051202912 | Not mapped | Not mapped | Not mapped | Not mapped | Not mapped | Not mapped |

## References

1. Meyer E, Kurian MA, Hayflick SJ. Neurodegeneration with Brain Iron Accumulation: Genetic Diversity and Pathophysiological Mechanisms. Annu Rev Genomics Hum Genet 16, 257–279 (2015).

2. Hayflick SJ, Kurian MA, Hogarth P. Neurodegeneration with brain iron accumulation. Handbook of clinical neurology 147, 293–305 (2018).

3. Hayflick SJ, et al. beta-Propeller protein-associated neurodegeneration: a new X-linked dominant disorder with brain iron accumulation. Brain 136, 1708–1717 (2013).

4. Haack TB, et al. Exome sequencing reveals de novo WDR45 mutations causing a phenotypically distinct, X-linked dominant form of NBIA. American journal of human genetics 91, 1144–1149 (2012).

5. Saitsu H, et al. De novo mutations in the autophagy gene WDR45 cause static encephalopathy of childhood with neurodegeneration in adulthood. Nature genetics 45, 445–449, 449e441 (2013).

6. Zhao YG, et al. The autophagy gene Wdr45/Wipi4 regulates learning and memory function and axonal homeostasis. Autophagy 11, 881–890 (2015).

7. Bakula D, et al. WIPI3 and WIPI4 beta-propellers are scaffolds for LKB1-AMPK-TSC signalling circuits in the control of autophagy. 8, 15637 (2017).

8. Proikas-Cezanne T, Takacs Z, Donnes P, Kohlbacher O. WIPI proteins: essential PtdIns3P effectors at the nascent autophagosome. Journal of cell science 128, 207–217 (2015).

9. Paudel R, et al. Neuropathology of Beta-propeller protein associated neurodegeneration (BPAN): a new tauopathy. Acta neuropathologica communications 3, 39 (2015).

10. Teinert J, Behne R, Wimmer M, Ebrahimi-Fakhari D. Novel insights into the clinical and molecular spectrum of congenital disorders of autophagy. Journal of inherited metabolic disease, (2019).

11. Choi AM, Ryter SW, Levine B. Autophagy in human health and disease. The New England journal of medicine 368, 651–662 (2013).

12. Stead ER, et al. Agephagy - Adapting Autophagy for Health During Aging. Frontiers in cell and developmental biology 7, 308 (2019).

13. Agrotis A, Ketteler R. On ATG4B as Drug Target for Treatment of Solid Tumours-The Knowns and the Unknowns. Cells 9, (2019).

14. Agrotis A, von Chamier L, Oliver H, Kiso K, Singh T, Ketteler R. Human ATG4 autophagy proteases counteract attachment of ubiquitin-like LC3/GABARAP proteins to other cellular proteins. The Journal of biological chemistry 294, 12610–12621 (2019).

15. Baskaran S, Ragusa MJ, Boura E, Hurley JH. Two-site recognition of phosphatidylinositol 3-phosphate by PROPPINs in autophagy. Molecular cell 47, 339–348 (2012).

16. Smith TF, Gaitatzes C, Saxena K, Neer EJ. The WD repeat: a common architecture for diverse functions. Trends Biochem Sci 24, 181–185 (1999).

17. Li D, Roberts R. WD-repeat proteins: structure characteristics, biological function, and their involvement in human diseases. Cell Mol Life Sci 58, 2085–2097 (2001).

18. Lu Q, et al. The WD40 repeat PtdIns(3)P-binding protein EPG-6 regulates progression of omegasomes to autophagosomes. Dev Cell 21, 343–357 (2011).

19. Obara K, Sekito T, Niimi K, Ohsumi Y. The Atg18-Atg2 complex is recruited to autophagic membranes via phosphatidylinositol 3-phosphate and exerts an essential function. The Journal of biological chemistry 283, 23972–23980 (2008).

20. Nakatogawa H, Suzuki K, Kamada Y, Ohsumi Y. Dynamics and diversity in autophagy mechanisms: lessons from yeast. Nature reviews Molecular cell biology 10, 458–467 (2009).

21. Wan H, et al. WDR45 contributes to neurodegeneration through regulation of ER homeostasis and neuronal death. Autophagy, 1–17 (2019).

22. Seibler P, et al. Iron overload is accompanied by mitochondrial and lysosomal dysfunction in WDR45 mutant cells. Brain, (2018).

23. Fusaki N, Ban H, Nishiyama A, Saeki K, Hasegawa M. Efficient induction of transgene-free human pluripotent stem cells using a vector based on Sendai virus, an RNA virus that does not integrate into the host genome. *Proceedings of the Japan Academy Series B*, Physical and biological sciences 85, 348–362 (2009).

24. Gasteiger E, Gattiker A, Hoogland C, Ivanyi I, Appel RD, Bairoch A. ExPASy: The proteomics server for in-depth protein knowledge and analysis. Nucleic acids research 31, 3784–3788 (2003).

25. Lenz M, et al. Epigenetic biomarker to support classification into pluripotent and non-pluripotent cells. Scientific reports 5, 8973 (2015).

26. Ng J, et al. Gene therapy restores dopamine transporter expression and ameliorates pathology in iPSC and mouse models of infantile parkinsonism. Science translational medicine 13, (2021).

27. Kirkeby A, et al. Generation of regionally specified neural progenitors and functional neurons from human embryonic stem cells under defined conditions. Cell reports 1, 703–714 (2012).

28. Tomoda K, et al. Derivation conditions impact X-inactivation status in female human induced pluripotent stem cells. Cell Stem Cell 11, 91–99 (2012).

29. Tchieu J, et al. Female human iPSCs retain an inactive X chromosome. Cell Stem Cell 7, 329–342 (2010).

30. Bar S, Seaton LR, Weissbein U, Eldar-Geva T, Benvenisty N. Global Characterization of X Chromosome Inactivation in Human Pluripotent Stem Cells. Cell reports 27, 20–29.e23 (2019).

31. Mekhoubad S, Bock C, de Boer AS, Kiskinis E, Meissner A, Eggan K. Erosion of dosage compensation impacts human iPSC disease modeling. Cell Stem Cell 10, 595–609 (2012).

32. Comertpay S, et al. Evaluation of clonal origin of malignant mesothelioma. J Transl Med 12, 301 (2014).

33. Love MI, Huber W, Anders S. Moderated estimation of fold change and dispersion for RNA-seq data with DESeq2. Genome biology 15, 550 (2014).

34. Mi H, Muruganujan A, Thomas PD. PANTHER in 2013: modeling the evolution of gene function, and other gene attributes, in the context of phylogenetic trees. Nucleic acids research 41, D377–386 (2013).

35. The Gene Ontology resource: enriching a GOld mine. Nucleic acids research 49, D325–d334 (2021).

36. Ashburner M, et al. Gene ontology: tool for the unification of biology. The Gene Ontology Consortium. Nature genetics 25, 25–29 (2000).

37. Kanehisa M, Furumichi M, Tanabe M, Sato Y, Morishima K. KEGG: new perspectives on genomes, pathways, diseases and drugs. Nucleic acids research 45, D353–d361 (2017).

38. Kanehisa M, Goto S, Furumichi M, Tanabe M, Hirakawa M. KEGG for representation and analysis of molecular networks involving diseases and drugs. Nucleic acids research 38, D355–360 (2010).

39. Kanehisa M, Sato Y, Kawashima M, Furumichi M, Tanabe M. KEGG as a reference resource for gene and protein annotation. Nucleic acids research 44, D457–462 (2016).

40. Jiao X, et al. DAVID-WS: a stateful web service to facilitate gene/protein list analysis. Bioinformatics 28, 1805–1806 (2012).

41. Mitre M, Mariga A, Chao MV. Neurotrophin signalling: novel insights into mechanisms and pathophysiology. Clin Sci (Lond) 131, 13–23 (2017).

42. Chen X, Yu C, Kang R, Tang D. Iron Metabolism in Ferroptosis. Front Cell Dev Biol 8, 590226 (2020).

43. Wu Q, Maniatis T. A striking organization of a large family of human neural cadherin-like cell adhesion genes. Cell 97, 779–790 (1999).

44. De Wolf V, et al. A complex Xp11.22 deletion in a patient with syndromic autism: exploration of FAM120C as a positional candidate gene for autism. American journal of medical genetics Part A **164a**, 3035–3041 (2014).

45. Nishimoto S, Kusakabe M, Nishida E. Requirement of the MEK5-ERK5 pathway for neural differentiation in Xenopus embryonic development. EMBO Rep 6, 1064–1069 (2005).

46. Zou J, et al. Targeted deletion of ERK5 MAP kinase in the developing nervous system impairs development of GABAergic interneurons in the main olfactory bulb and behavioral discrimination between structurally similar odorants. The Journal of neuroscience : the official journal of the Society for Neuroscience 32, 4118–4132 (2012).

47. Hetz C, Papa FR. The Unfolded Protein Response and Cell Fate Control. Molecular cell 69, 169–181 (2018).

48. Brunetti-Pierri N, Scaglia F. GM1 gangliosidosis: review of clinical, molecular, and therapeutic aspects. Molecular genetics and metabolism 94, 391–396 (2008).

49. Mohammad SS, et al. Magnetic resonance imaging pattern recognition in childhood bilateral basal ganglia disorders. Brain Commun 2, fcaa178 (2020).

50. Regier DS, et al. MRI/MRS as a surrogate marker for clinical progression in GM1 gangliosidosis. American journal of medical genetics Part A 170, 634–644 (2016).

51. Kukkonen JP. Orexin/Hypocretin Signaling. Curr Top Behav Neurosci 33, 17–50 (2017).

52. Wilson JL, et al. Consensus clinical management guideline for beta-propeller protein-associated neurodegeneration. Developmental medicine and child neurology, (2021).

53. Lee JR. Protein tyrosine phosphatase PTPRT as a regulator of synaptic formation and neuronal development. BMB Rep 48, 249–255 (2015).

54. Nam J, Mah W, Kim E. The SALM/Lrfn family of leucine-rich repeat-containing cell adhesion molecules. Semin Cell Dev Biol 22, 492–498 (2011).

55. Pei YP, et al. ICAM5 as a Novel Target for Treating Cognitive Impairment in Fragile X Syndrome. The Journal of neuroscience : the official journal of the Society for Neuroscience 40, 1355–1365 (2020).

56. Fernandez RF, et al. Acyl-CoA synthetase 6 enriches the neuroprotective omega-3 fatty acid DHA in the brain. Proceedings of the National Academy of Sciences of the United States of America 115, 12525–12530 (2018).

57. Gottlieb RA, Andres AM, Sin J, Taylor DP. Untangling autophagy measurements: all fluxed up. Circ Res 116, 504–514 (2015).

58. Yoshii SR, Mizushima N. Monitoring and Measuring Autophagy. International journal of molecular sciences 18, (2017).

59. Mauthe M, et al. Chloroquine inhibits autophagic flux by decreasing autophagosome-lysosome fusion. Autophagy 14, 1435–1455 (2018).

60. Pelz O, Gilsdorf M, Boutros M. web cellHTS2: a web-application for the analysis of high-throughput screening data. BMC Bioinformatics 11, 185 (2010).

61. Waguri S, Komatsu M. Biochemical and morphological detection of inclusion bodies in autophagy-deficient mice. Methods Enzymol 453, 181–196 (2009).

62. Kraja AT, et al. Associations of Mitochondrial and Nuclear Mitochondrial Variants and Genes with Seven Metabolic Traits. American journal of human genetics 104, 112–138 (2019).

63. Liang C, et al. Autophagic and tumour suppressor activity of a novel Beclin1-binding protein UVRAG. Nature cell biology 8, 688–699 (2006).

64. Agrotis A, Pengo N, Burden JJ, Ketteler R. Redundancy of human ATG4 protease isoforms in autophagy and LC3/GABARAP processing revealed in cells. Autophagy 15, 976–997 (2019).

65. Barral S, Kurian MA. Utility of Induced Pluripotent Stem Cells for the Study and Treatment of Genetic Diseases: Focus on Childhood Neurological Disorders. Frontiers in molecular neuroscience 9, 78 (2016).

66. Xiong Q, et al. WDR45 Mutation Impairs the Autophagic Degradation of Transferrin Receptor and Promotes Ferroptosis. Front Mol Biosci 8, 645831 (2021).

67. Fu XH, et al. COL1A1 affects apoptosis by regulating oxidative stress and autophagy in bovine cumulus cells. Theriogenology 139, 81–89 (2019).

68. Paiva I, et al. Alpha-synuclein deregulates the expression of COL4A2 and impairs ER-Golgi function. Neurobiology of disease 119, 121–135 (2018).

69. Tang ME, et al. Matrix metalloproteinase-degraded type I collagen is associated with APOE/TOMM40 variants and preclinical dementia. Neurology Genetics 6, e508 (2020).

70. Cescon M, Chen P, Castagnaro S, Gregorio I, Bonaldo P. Lack of collagen VI promotes neurodegeneration by impairing autophagy and inducing apoptosis during aging. Aging (Albany NY) 8, 1083–1101 (2016).

71. Stanga D, Zhao Q, Milev MP, Saint-Dic D, Jimenez-Mallebrera C, Sacher M. TRAPPC11 functions in autophagy by recruiting ATG2B-WIPI4/WDR45 to preautophagosomal membranes. *Traffic (Copenhagen*, Denmark) 20, 325–345 (2019).

72. Chang CY, et al. Induced Pluripotent Stem Cell (iPSC)-Based Neurodegenerative Disease Models for Phenotype Recapitulation and Drug Screening. Molecules 25, (2020).

73. Garcia-Leon JA, Vitorica J, Gutierrez A. Use of human pluripotent stem cell-derived cells for neurodegenerative disease modeling and drug screening platform. Future Med Chem 11, 1305–1322 (2019).

74. Little D, Ketteler R, Gissen P, Devine MJ. Using stem cell-derived neurons in drug screening for neurological diseases. Neurobiol Aging 78, 130–141 (2019).

75. Papandreou A, Luft C, Barral S, Kriston-Vizi J, Kurian MA, Ketteler R. Automated high-content imaging in iPSC-derived neuronal progenitors. SLAS Discov 28, 42–51 (2023).

76. Celsi F, et al. Mitochondria, calcium and cell death: a deadly triad in neurodegeneration. Biochimica et biophysica acta 1787, 335–344 (2009).

77. Guo T, Zhang D, Zeng Y, Huang TY, Xu H, Zhao Y. Molecular and cellular mechanisms underlying the pathogenesis of Alzheimer’s disease. Molecular neurodegeneration 15, 40 (2020).

78. Moore DJ, West AB, Dawson VL, Dawson TM. Molecular pathophysiology of Parkinson’s disease. Annual review of neuroscience 28, 57–87 (2005).

79. Hansen TE, Johansen T. Following autophagy step by step. BMC Biol 9, 39 (2011).

80. Hundeshagen P, Hamacher-Brady A, Eils R, Brady NR. Concurrent detection of autolysosome formation and lysosomal degradation by flow cytometry in a high-content screen for inducers of autophagy. BMC Biol 9, 38 (2011).

81. Liu Y, Levine B. Autosis and autophagic cell death: the dark side of autophagy. Cell Death Differ 22, 367–376 (2015).

82. Wang Y, et al. Cardiac glycosides induce autophagy in human non-small cell lung cancer cells through regulation of dual signaling pathways. Int J Biochem Cell Biol 44, 1813–1824 (2012).

83. Dunn DE, He DN, Yang P, Johansen M, Newman RA, Lo DC. In vitro and in vivo neuroprotective activity of the cardiac glycoside oleandrin from Nerium oleander in brain slice-based stroke models. J Neurochem 119, 805–814 (2011).

84. Wang JKT, et al. Cardiac glycosides provide neuroprotection against ischemic stroke: discovery by a brain slice-based compound screening platform. Proceedings of the National Academy of Sciences of the United States of America 103, 10461–10466 (2006).

85. Elmaci İ, Alturfan EE, Cengiz S, Ozpinar A, Altinoz MA. Neuroprotective and tumoricidal activities of cardiac glycosides. Could oleandrin be a new weapon against stroke and glioblastoma? Int J Neurosci 128, 865–877 (2018).

86. Rossignoli G, et al. Aromatic l-amino acid decarboxylase deficiency: a patient-derived neuronal model for precision therapies. Brain 144, 2443–2456 (2021).

87. Kirkeby A, Nelander J, Parmar M. Generating regionalized neuronal cells from pluripotency, a step-by-step protocol. Frontiers in cellular neuroscience 6, 64 (2012).

88. Huang da W, Sherman BT, Lempicki RA. Systematic and integrative analysis of large gene lists using DAVID bioinformatics resources. Nature protocols 4, 44–57 (2009).

89. Mi H, et al. PANTHER version 16: a revised family classification, tree-based classification tool, enhancer regions and extensive API. Nucleic acids research 49, D394–d403 (2021).

90. Bindea G, et al. ClueGO: a Cytoscape plug-in to decipher functionally grouped gene ontology and pathway annotation networks. Bioinformatics 25, 1091–1093 (2009).

